# A Novel Factor in Olfactory Ensheathing Cell-Astrocyte Crosstalk: Anti-Inflammatory Protein α-Crystallin B

**DOI:** 10.1101/2020.08.28.273003

**Authors:** Aybike Saglam, Anne L. Calof, Susan Wray

## Abstract

Astrocytes are key players in CNS neuroinflammation and neuroregeneration that may help or hinder recovery, depending on the context of the injury. Although pro-inflammatory factors that promote astrocyte-mediated neurotoxicity have been shown to be secreted by reactive microglia, anti-inflammatory factors that suppress astrocyte activation are not well-characterized. Olfactory ensheathing cells (OECs), glial cells that wrap axons of olfactory sensory neurons, have been shown to moderate astrocyte reactivity, creating an environment conducive to regeneration. Similarly, astrocytes cultured in medium conditioned by cultured OECs (OEC-CM) show reduced nuclear translocation of Nuclear Factor kappa-B (NFκB), a pro-inflammatory protein that induces neurotoxic reactivity in astrocytes. In this study, we screened primary and immortalized OEC lines to identify these factors and discovered that Alpha B-crystallin (CryAB), an antiinflammatory protein, is secreted by OECs via exosomes, coordinating an intercellular immune response. Our results showed: 1) OEC exosomes block nuclear NFκB translocation in astrocytes while exosomes from *CryAB*-null OECs could not; 2) OEC exosomes could be taken up by astrocytes and 3) CryAB treatment suppressed multiple neurotoxicity-associated astrocyte transcripts. Our results indicate that OEC-secreted factors are potential agents that can ameliorate, or even reverse, the growth-inhibitory environment created by neurotoxic reactive astrocytes following CNS injuries.

**Main Points:**

- Astrocytes uptake OEC-secreted exosomes.
- WT OEC-exosomes, but not CryAB-null OEC-exosomes, block nuclear NFκB translocation in astrocytes.
- CryAB, and other factors secreted by OECs, suppresses multiple neurotoxicity-associated astrocyte transcripts.

## 1 Introduction

Damage to the central nervous system (CNS) provokes morphological and molecular changes in astrocytes, causing them to become ‘reactive astrocytes’ (Liddelow & Barres, 2017). These reactive cells play positive roles during CNS injury, such as confining inflammation by surrounding the damaged tissue and creating a barrier between it and uninjured tissues (Silver, *et al*., 2015). Reactive astrocytes have been traditionally characterized by increased expression of intermediate filament proteins such as GFAP (Glial Fibrillary Acidic Protein), vimentin, and nestin (summarized in Liddelow & Barres, 2017). Excessive or sustained astrocyte reactivity is characterized by activation of pro-inflammatory pathways such as the NFκB pathway (Liddelow & Barres., 2017; Wheeler 2020). This activity can be deleterious to functional recovery, since it can lead to chronic inflammation and neurotoxicity (Sofroniew, *et al*., 2010). A better understanding of the molecular mechanisms that govern astrocyte reactivity would therefore be helpful to create environments conducive to regeneration following CNS injury.

The mammalian olfactory system shows robust neurogenesis throughout life. Data suggest that both neural niche signals and the surrounding glia, including olfactory ensheathing cells (OECs), give the olfactory mucosa this unique capability (Li, *et al*., 2005; Roet & Verhaagen, 2014). Several groups have transplanted OECs into CNS injury sites, and observed improved axonal regeneration (Li *et al*., 1997; Imaizumi *et al*., 2000), functional recovery (Johansson *et al*., 2005), reduced astrocytic scar tissue (Ramer *et al*., 2004), and an attenuated hostile astrocyte response (Lakatos *et al*, 2003; summarized in Roet & Verhaagen, 2014). Moreover, factors secreted by OECs have been shown to moderate astrocyte reactivity, at least insofar as their presence results in reduction of GFAP expression and nuclear translocation of NFκB (Chuah *et al*., 2011; O’Toole *et al*., 2007). Identification of molecules secreted by OECs, which specifically affect astrocyte reactivity, should lead not only to a better understanding of the crosstalk between astrocytes and OECs; it may also reveal mechanisms that can block the metamorphosis of astrocytes into neurotoxic cells.

To identify such molecules, this study used lipopolysaccharide (LPS)-treated astrocytes, a model for neurotoxic reactive astrocytes, and assessed conditioned medium (CM) from immortalized clonal mouse OEC cell lines (Calof & Guevara, 1993). Nuclear NFκB translocation in astrocytes was measured to determine if CM from these cell lines could mimic primary OEC-CM, which blocked the LPS pro-inflammatory response in astrocytes. Two immortalized cell lines were chosen for further study: one whose CM mimicked the effect of primary OECs (positive control); and a second, whose CM did not block the LPS response in astrocytes (negative control). These two cell lines and primary OECs were challenged with LPS, and the conditioned media screened by mass spectrometry. Using this strategy, the heat-shock protein CryAB (D’Agostino *et al*., 2013) was identified. Subsequent experiments showed that: 1) CryAB is secreted by OECs via exosomes; 2) exogenous CryAB suppressed LPS-induced astrocyte reactivity; 3) exosomes containing CryAB are taken up by astrocytes; and 4) unlike wildtype (WT) OEC-exosomes, *CryAB-null* (*CryAB*^−/−^) OEC exosomes fail to suppress LPS-induced astrocyte reactivity measured by nuclear NFκB translocation. Finally, examination of transcripts that are associated with neurotoxic-reactive astrocytes (Liddelow *et al*., 2017) revealed that either exogenous CryAB or OEC-CM can suppress expression of several of these transcripts. Taken together, the data indicate that CryAB secreted by OECs, via exosomes, is an important factor for OEC-astrocyte crosstalk that can block astrocytes from becoming neurotoxic cells. Ultimately, mimicking appropriate astrocyte-OEC crosstalk *in vivo* may contribute to an environment conducive to regeneration following a broad range of CNS injuries.

## 2 Materials & Methods

### 2.1 Mice

All mice were maintained, and all animal handling procedures were performed according to protocols approved by the National Institutes of Health NINDS Institutional Animal Care and Use Committee. *CryAB*-null (*CryAB^Del(9Hspb2-Cryab)1Wawr^*, henceforth referred to as *CryAB*^−/−^) mice (Brady *et al*., 2001) were obtained as homozygous sperm, revived by IVF using eggs from C57bl6/N mice (Jackson Laboratory), and resulting heterozygotes intercrossed to obtain obtain *CryAB*^−/−^ and *CryAB*^+/+^ (wildtype, WT) lines, which were used as the source for OEC primary cultures (see below). Mice were genotyped using a three-primer PCR protocol: 5’-TAGCTTAATAATCTGGGCCA-3’, 5’-GGAGTTCCACAGGAAGT-ACC-3’, and 5’-TGGAAGGATTGGAG-CTACGG-3’ primers were used in 4:1:1 molar ratio. Amplification was performed for 40 cycles at 94°C for 15 sec, 62°C for 30 sec and 72°C at 1 min. PCR produced a 310-bp product for the WT allele and a 600-bp product for the null allele.

### 2.2 Cell culture and reagents

Primary cultures of OECs were generated as described previously (Dairaghi *et al*., 2018). Briefly, olfactory bulbs of postnatal (PN) day 0-7 mice were collected and placed in an enzyme mix: 300μg/ml hyaluronidase (Sigma, Cat# H3631, St. Louis, MO), 30U/ml dispase I (Sigma, Cat# D4818), 1.2 mg/ml collagenase type 4 (Worthington, Cat# 43E14231, Lakewood, NJ), 10U/ml DNAse I (Worthington, Cat# 54E7315); for 35 min at 37°C with constant agitation (Au & Roskams, 2003). Cells were run through a 40μm cell strainer to remove non-dissociated tissue pieces and then washed with DMEM-F12 medium. Subsequently, cells were purified by the differential cell adhesion method (Nash *et al*., 2001), which consists of three steps: 1) Cells were seeded into uncoated T75 flasks (4×10^6^ viable cells/flask, VWR, Cat# 734-2788, Radnor, PA) for 18 hrs to remove fibroblasts; 2) the supernatant of the first step was seeded into another uncoated flask for up to 36 hrs to remove astrocytes; and finally 3) the supernatant of the second flask was seeded onto poly-L-lysine (Sigma, Cat# P4707)-coated flasks to grow primary OECs. Cells were cultured for up to 2 weeks and medium was changed every 2–3 days. OECs constituted more than 90% of the cells in the culture based on p75 and S100B immunostaining (data not shown). For OECs to be co-cultured with primary astrocytes, the medium was gradually changed to serum-free medium (Klenke & Taylor-Burds, 2012) supplemented with 5ng/ml HB-EGF (PeproTech, Cat# 100-47, Rocky Hill, NJ), and B27 (Thermo Fisher Scientific, Cat# A3582901) to provide a medium compatible with astrocyte culture, since serum has been shown to induce astrocyte reactivity (Foo *et al*., 2011).

Primary astrocyte cultures were obtained by magnetic sorting as previously described (Holt *et al*., 2019), with some modifications. Briefly, 10-20 cortices of PN day 2-4 pups were dissociated using the MACS Neural Tissue Dissociation Kit-T (Miltenyi Biotec, Cat# 130-093-231, Auburn, CA) at 37°C (5% CO^2^, 30 min). Non-dissociated tissue was removed using a 40μm cell strainer (Fisher Scientific, Cat# 22-363-547), and the remaining cell solution was centrifuged (300g, 5 min). Next, a discontinuous density gradient, prepared using 1:1 albumin-ovomucoid solution (10mg/ml of each) (Worthington, Cat# OI; GeminiBio, Cat# 700-102P, West Sacramento, CA), was used to remove cell debris and inhibit enzyme activity. The cell pellet was resuspended in 80μl Hank’s Balanced Salt Solution (HBSS) (Gibco, Cat# 14025-092) plus 20μl anti-GLAST (ACSA-1) MicroBeads (Miltenyi Biotec, Cat# 130-095-825, Auburn, CA) for up to 10^7^ cells, and incubated for 10 min (4°C). Cells were washed and incubated in 90μl HBSS plus 10μl anti-Biotin MicroBeads for another 15 min (4°C) before running through MACS column for positive selection of astrocytes. Cells were cultured for one week and then the same procedure was followed with anti-Prominin-1 MicroBeads (Miltenyi Biotec, Cat# 130-092-564) for the negative selection of radial glia, followed by another positive selection with anti-GLAST antibody the same day, to increase the purity of astrocyte cultures. Sorted cells were cultured in T25 flasks coated with poly-L-lysine, in 5ml serum-free astrocyte culture medium (ACM, described above). In our hands, astrocytes isolated by this method and cultured in ACM were not reactive when stained with NFκB (not shown). The same method was adjusted to obtain oligodendrocyte cultures using anti-O4-MicroBeads (Miltenyi Biotec, Cat# 130-096-670). The immortalized mouse astrocyte line C8D30 (ATCC, VA, USA) was cultured in DMEM-F12 (Gibco, Cat# 10313-02, 11765-054, Long Island, NY) containing 10% Fetal Bovine Serum (FBS) (Gibco, Cat# 10438-026), plus 0.5% antibiotic-antimycotic (Gibco, Cat# 15240-062) at 37°C in 5%CO2.

### 2.3 Immortalized OEmyc790 Cell Lines

Six immortalized OEC lines (OEmyc790-C7s.D, D6s.AB8, C6s.BG9, D10 and D4), derived from retrovirus-mediated transformation of primary embryonic mouse olfactory epithelium cultures derived from E15 mouse embryos (Calof & Guevara, 1993), were analyzed; two lines, OEmyc790-C7s.D (C7) and OEmyc790-C4 (C4), were used for the studies described below. Cells were plated on cell culture plates (Fisher Scientific, Cat# 130190, Waltham, MA) and cultured in DMEM-F12 as described above. Medium was changed every 3-5 days. When 60% confluent, a 1:4 dilution of trypsin was used (Gibco, Cat# 15400054) to split the cells into thirds.

### 2.4 Primary antibodies and recombinant proteins

The following antibodies were used: CryAB rabbit polyclonal antibody (Millipore, Cat# ABN185, Darmstadt, Germany, 1:1K for WB, 1:4K for immunofluorescence (IF); Histone mouse monoclonal antibody (Fisher Scientific, Cat# AHO1432, Waltham, MA, 1:200 for WB); NFκB rabbit polyclonal antibody (C-20, Santa Cruz, Cat# sc-372, Santa Cruz, CA, 1:650 for WB, 1:750 for IF); Sox10 goat polyclonal antibody (N-20, Santa Cruz, Cat# sc-17342, 1:300 for IF); Alix mouse monoclonal antibody (3A9, Cell Signaling, Cat# 2171T, 1:1K for WB); GFAP chicken polyclonal antibody (Aves, 1:4K for IF); Flotillin-1 rabbit polyclonal antibody (D2V7J, Cell Signaling, Cat# 18634, 1:1K for WB); β-actin mouse monoclonal antibody (AC-74, Millipore, Cat# A2228, 1:1K for WB); Tomm20 rabbit polyclonal antibody (FL-145, Santa Cruz, Cat# sc-11415, 1:1K for WB); CD63 biotinylated antibody (Miltenyi Biotec, Cat# 130-108-922, Auburn, CA, 1:15 for IF); BLBP mouse monoclonal antibody (Abcam, Cat# ab131137, 1:2K for IF) and p75-NGFR rabbit polyclonal antibody (Millipore, Cat# AB1554, 1:5K). Recombinant chicken Anosmin1 (MyBioSource, Cat# MBS963562-COA, San Diego, CA) was used at 5nM while recombinant CryAB protein (MyBioSource, Cat# MBS964495) and recombinant myoglobin (MyBioSource, Cat# MBS142891) were used at 50ng/ml unless stated otherwise.

### 2.5 Mass spectrometry

Primary OECs, and immortalized C7 and C4 OEC cell lines, were established by seeding them in T75 flasks at a concentration of 8×10^5^ cells/flask in regular growth medium. To concentrate secreted proteins, the cells in each flask were rinsed and media replaced with 10ml/flask of Earle’s Balanced Salt Solution (EBSS) with 5.5mM D-Glucose (Gibco, Cat# 14155-063); conditioned medium (CM) was collected after 48 hrs total of incubation. For the last 2 hrs of the 48-hr collection period, either 1μl/ml LPS (Sigma, Cat# L6529) or 5nM recombinant chicken Anosmin1 was added. CM was then collected, centrifuged to remove debris, and frozen at −80°C. Frozen samples were freeze-dried using a lyophilizer (Novalyphe-NL150, Savant Instruments, Holbrook, NY). The pellets were reconstituted in water and bicinchoninic acid (BCA) protein assay was performed. 200μg/60ul protein per group was submitted for mass spectrometry analysis (NINDS Protein Facility, NIH). Each sample was digested with trypsin. Tandem Mass Tag (TMT) labeled samples were mixed together (TMT 126-131). The mixture was separated using hydrophilic interaction liquid chromatography (HILIC) high performance liquid chromatography (HPLC) system. Five HILIC fractions were collected from the mixed sample. One liquid chromatography-tandem mass spectrometry (LC/MS/MS) experiment was performed for each HILIC fraction, using an Orbitrap Fusion Lumos Mass Spectrometer (Thermo Fisher Scientific, Waltham, MA) coupled to a 3000 Ultimate high-pressure liquid chromatography instrument (Thermo Fisher Scientific). Proteome Discoverer 2.2 software used for database search and TMT quantification, and data were mapped against the Sprot mouse database. “Primary OEC+LPS-CM” was used as reference to calculate the ratio for LPS treated samples; “Primary OEC+5nMA1 −CM” was used as reference to calculate the ratio for 5nMAnosmin1-treated samples. No normalization was performed. See Supplemental Data 1 and 2 for obtained values.

### 2.6 Immunoblot analysis

As a readout of reactivity, quantitative immunoblot analysis was performed on nuclear fractions of immortalized C8D30 astrocytes treated with 1μl/ml LPS or vehicle control for 2 hrs. For co-culture groups, OECs seeded on porous inserts (0.4μm Millicell Cell Culture Insert, Millipore, Cat# PICM0RG50) were placed on top of astrocytes for 24 hrs and were discarded at the end of the incubation period, so that only astrocytes were collected for subsequent protein analysis. For the CM treated groups, CM from each line was collected (after 24 hr incubation) and then added to astrocytes for 22 hrs, followed by a 2-hour LPS treatment. Astrocytes were then collected by scraping and the CNMCS Compartmental Protein Extraction Kit (BioChain Cat# K3013010 Hayward, CA) with protease/phosphatase inhibitors (PI, Cell Signaling, Cat# 5872S, Danvers, MA) was used and the nuclear fractions were isolated for each treatment condition. The fractions were run on BioRad Mini-Protean TGX Stain-Free Gels (Cat#4568084), transferred to PVDF stain-free blot (Trans-Blot Turbo Transfer Pack, Cat#1704156) via the Trans-Blot Turbo transfer system (BioRad), and blocked with 5% dry milk (BioRad, Cat #170–6404) prior to staining with NFκB antibody. Membranes were exposed to Clarity enhanced chemiluminescence (ECL) reagent (Cat. # 170–5061, Bio-Rad) for 5 min and the signal was detected using ChemiDoc MP (Cat. # 170–8280, Bio-Rad). Quantification of band intensities was calculated using Image Lab 5.0 software (Bio-Rad) and normalized by the loading control immunostained for Histone on the same sample and the same blot. Three biological replicates were used for statistical analysis.

### 2.7 Quantitative immunofluorescence

Fluorescent immunostaining for nuclear NFκB and cytoplasmic NFκB was quantified in immortalized C8D30 astrocytes following 2-hr treatment with 1μg/ml LPS or a cocktail of 3 cytokines: Il-1α (3ng/ml, Sigma, Cat# I3901), TNFα (30ng/ml, Cell Signaling, Cat# 8902SF) and C1q (400ng/ml, MyBioSource, Cat# MBS143105, San Diego, CA), as follows: After immunofluorescence staining for NFκB, confocal images were taken on a Zeiss LSM 800 Confocal Microscope (Carl Zeiss, Thornwood, NY). A defined area was measured in both nuclear and cytoplasmic compartments for each astrocyte, and the fluorescence intensity was quantified for each area in each cell using Imaris software. The ratio of nuclear to cytoplasmic fluorescence intensity was used as a quantitative readout of astrocyte reactivity. Median values were calculated for each biological replicate (N=3) obtained from multiple images (2-3/well) containing a total of ~100 cells/treatment (Figure 1C, cell numbers in Supplemental Table 1) or ~50 cells/treatment (Figure 2B, cell numbers in Supplemental Table 2). Values greater than one indicate that the NFκB value was higher in the nucleus compared to cytoplasm, and cells were reactive. Statistics (ANOVA) were performed and the average median value ±standard deviation (SD) per treatment plotted.

**Figure 1:**
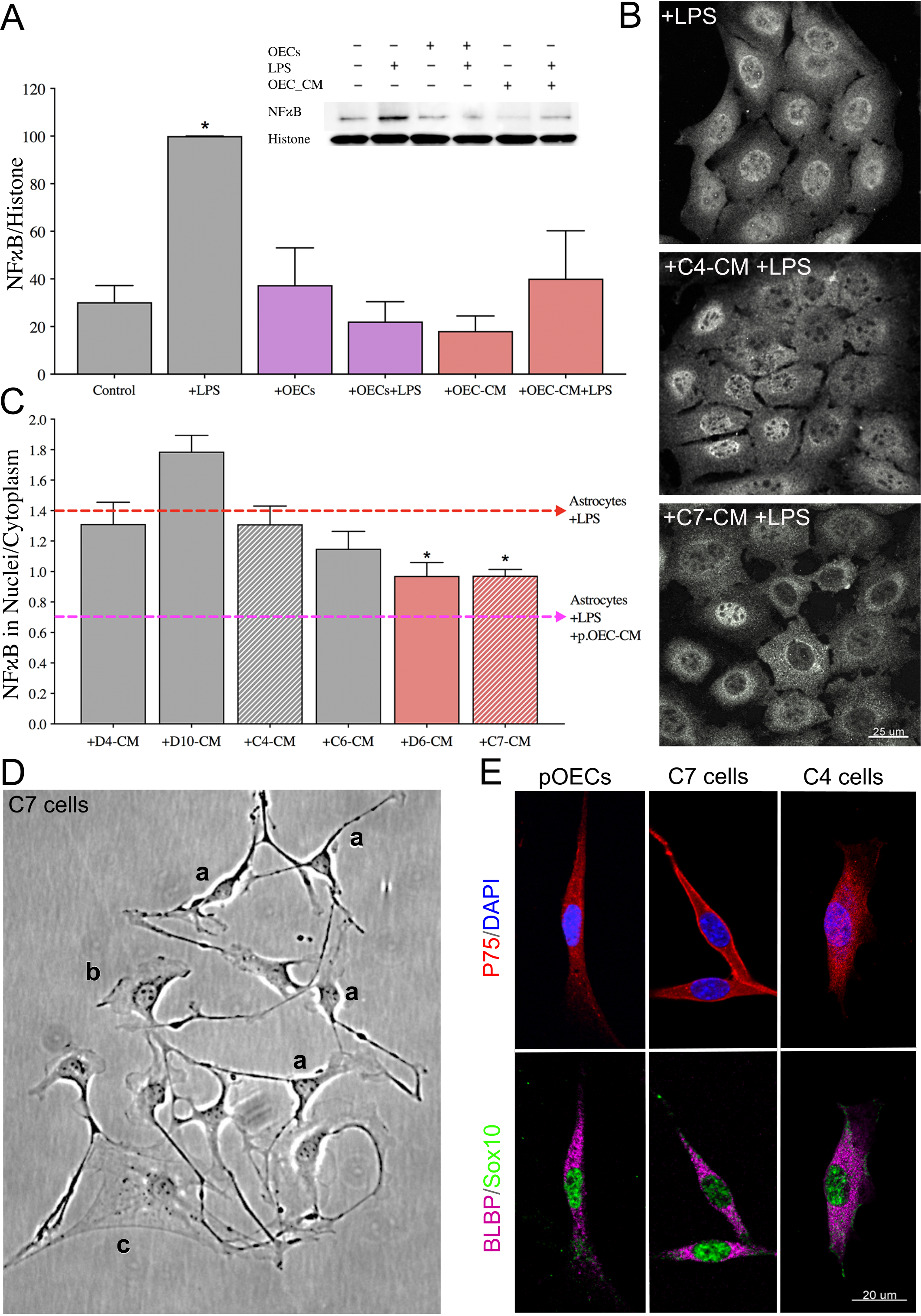
OEC-CM alone is sufficient to suppress LPS-induced astrocyte reactivity, an effect mimicked by a subset of immortalized OEC lines. (A) Quantitative immunoblotting for NFκB was performed on the nuclear fraction of C8D30 astrocyte lysates (inset). All groups were compared using a one-way ANOVA (N=3). Treatment with LPS for 2 hrs significantly increased NFκB activity (* indicates p≤0.05, gray bars). The presence of OECs blocked the effect of LPS (purple bars). OEC conditioned medium (OEC-CM) alone also blocked the increase in nuclear NFκB (red bars). (B and C) CM from six immortalized OEC lines were investigated for their ability to block the effect of LPS on nuclear NFκB translocation in C8D30 astrocytes, as measured by the ratio of fluorescence intensity of nuclear to cytoplasmic NFκB. (B) Photomicrograph of images of C8D30 astrocytes cocultured with CM of immortalized cell lines. (C) Fluorescent NFκB nuclear and cytoplasmic intensities were measured and ratios plotted. Pink dashed line depicts value of astrocytes treated with primary OEC-CM +LPS and red dashed line depicts value of astrocytes +LPS. A median value was calculated from ~100 cells per field, and then a mean/group was calculated. C7-CM and D6-CM decreased NFκB nuclear translocation (N = 3; * p ≤ 0.05; one-way ANOVA), while C4-CM treated groups was not significantly different than astrocytes +LPS alone (N = 3; * p<0.05; one-way ANOVA). (D) Photomicrograph from line C7. Multiple morphologies were found in all the OEC cell lines: (a) Schwann Cell-like, (b) astrocyte-like type1, and (c) astrocyte-like type2. Schwann cell-like spindle cells (a) predominated in C7. (E) C7 and C4 olfactory cell lines share multiple markers with OECs including BLBP (magenta), Sox10 (green) and P75 (red). Scale bars represent 25 and 20μm.

**Figure 2:**
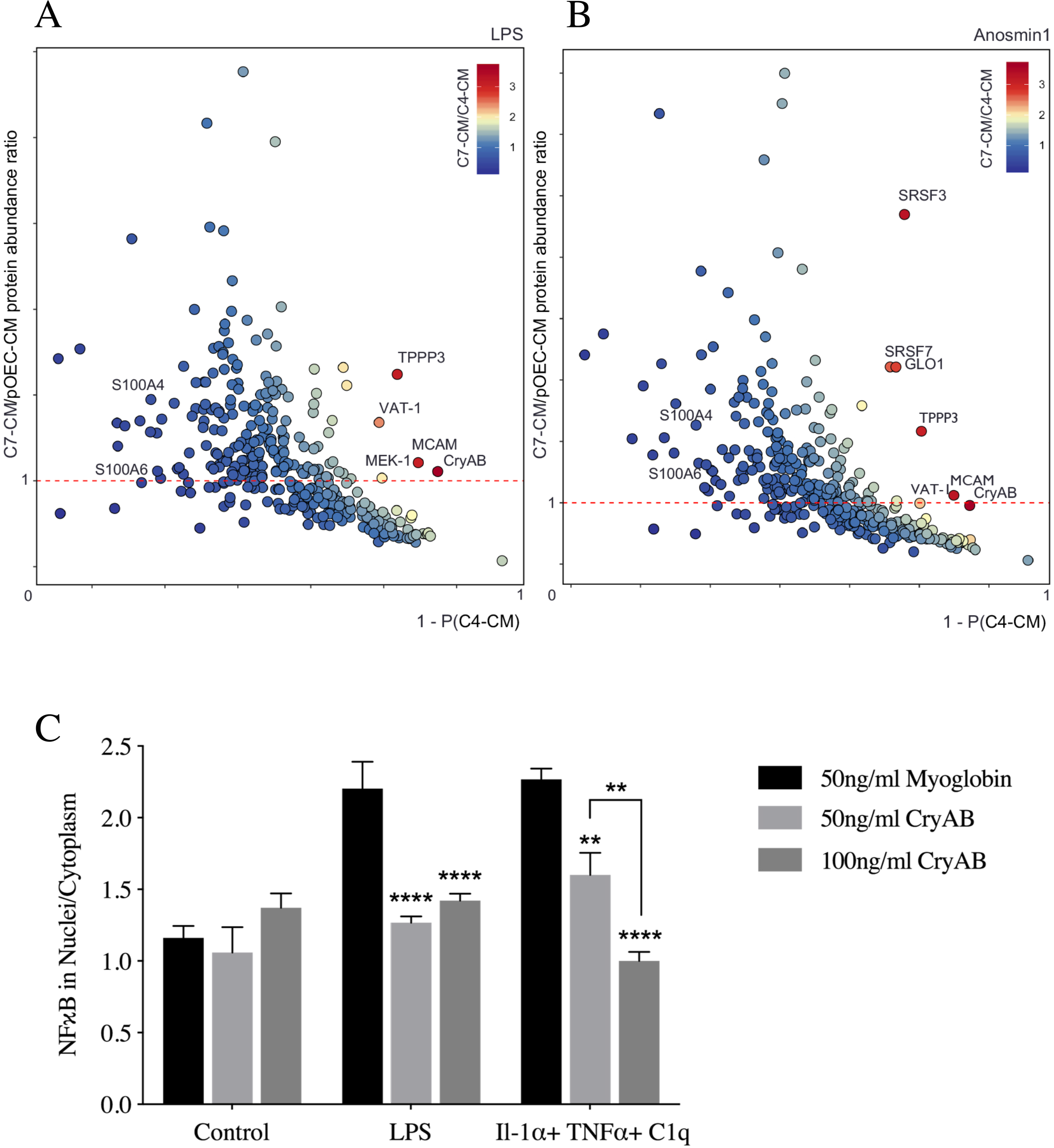
OEC-secreted anti-inflammatory protein CryAB and recombinant CryAB is sufficient to suppress astrocyte reactivity measured by NFκB. (A, B) Comparison of factors secreted from C7 line, C4 line and primary OECs (pOECs), analyzed by mass spectrometry. Proteins detected in CM following LPS treatment (A) or Anosmin1 treatment (B) were ranked by their relative abundance indicated by color code (heat map, inset). Relative abundance of detected proteins in C7-CM compared to pOEC-CM is graphed on the Y axes, and relative absence of the same proteins from C4-CM (1-P(C4-CM) = probability of not being found in C4-CM) is graphed on the X axes. Proteins of similar abundance in CMs from C7 cells and pOECs (horizontal red-dashed lines), and not likely to be present in C4-CM (Y axes) were identified. Alpha crystallin B chain (CryAB) had the highest C7/C4 expression ratio and was equally abundant in CM from C7 cells and pOECs. (C) Recombinant CryAB alone suppressed the inflammatory response, quantified as the ratio of nuclear to cytoplasmic NFκB (Y axis) in astrocytes exposed to either LPS or a cocktail of the cytokines Il-1α, TNFα and C1q for 2 hrs. Group values were obtained from triplicate wells in which a median value was calculated from ~50 cells per field. N = 3; (*p ≤ 0.05; **p ≤ 0.01; ****p ≤ 0.0001;) two-way ANOVA.

### 2.8 Isolation of exosomes and exosome uptake experiments

For isolation of exosomes, the protocol of Adolf and colleagues (2018) was used with slight modifications. Briefly, 24 hr prior to exosome collection, cells were washed and medium changed to exosome depleted medium (EDM) containing 10% exosome-depleted FBS (Gibco, Cat# A27208-03). CM was collected, (protease inhibitor (PI) was added immediately for immunoblotting) and samples kept at 4°C until exosome isolation. Exosomes were isolated through three centrifugation steps: 1) CM was spun for 10 min at 2,000*g* to remove debris; 2) the resulting supernatant was centrifuged for 30 min at 10,000*g* to pellet microvesicles; and 3) this second supernatant was centrifuged for 4 hrs at 100,000*g* (Optima MAX-XP ultracentrifuge, TLA-100.3 rotor, Beckman Coulter). Following ultracentrifugation, pelleted exosomes were re-suspended in buffer (for ELISA and immunoblotting) or cell culture medium, as required. For astrocyte uptake experiments, isolated OEC-exosomes were resuspended by pipetting and added directly to the culture medium of *CryAB*^−/−^ astrocytes for 4 hrs. Cultures were then fixed with 4% paraformaldehyde and stained for markers of interest. Images were taken on a stimulated emission depletion (STED) confocal microscope (Leica, Wetzlar, Germany) for the visualization of internalized exosomes in astrocytes.

### 2.9 CryAB Immunoprecipitation

CryAB was immunoprecipitated (IP-CryAB) from isolated OEC-exosome fractions that were lysed in RIPA buffer. Briefly, 200μl Protein A Dynabeads (30mg/ml, Invitrogen, Cat# 10001D, Carlsbad, CA) were washed 3 times in PBS+ 0.1% Tween (PBST) using a magnetic stand (Millipore, PureProteome Cat# LSKMAGS08), CryAB antibody (400μl, 1:50 (10 μg/mL) in PBST) was added to the beads, and the mixture was incubated (30 min, RT) with constant agitation. The antibody solution was removed, beads washed (3x), exosome fractions resuspended in PBS were added, and the mixture was incubated overnight (4°C). Beads were then washed (4x) and CryAB protein eluted by addition of 60μl of 0.2M Glycine (pH 2.5); the pH of the eluate was neutralized by addition of 5μl of 1M Tris (pH 8.5). Cell culture, immunoblotting or ELISA was performed.

### 2.10 ELISA

CryAB concentration was measured in exosome fractions using a competitive ELISA kit (MyBioSource, Cat# MBS7239470, San Diego, CA) according to manufacturer’s instructions. Isolated exosomes were sonicated and lysed in Buffer M (containing NP40) plus PI from Protein Extraction Kit (BioChain). Equivalent quantities of total exosomal protein or supernatant CM protein, determined by BCA protein assay, were brought to equivalent volumes in EDM. 100μl of samples were added to each well and measured with a microplate reader (FlexStation 3; Molecular Devices, Sunnyvale, CA). Results were analyzed with SoftMax Pro Software (Molecular Devices).

### 2.11 Quantitative RT-PCR (q-RT-PCR)

cDNA synthesis was performed using Superscript™ III reverse transcriptase (Invitrogen), and PCR carried out using the ViiA7 Real-Time PCR System (Applied Biosystems, Waltham, MA) in 20μl final volume, containing 10μl of SsoAdvanced Universal SYBR Green Supermix (BioRad Cat#1725271), 2μl of a primer mix with a concentration of 1μM of each primer and 1μl of cDNA and 7μl water. Samples were run in triplicate. The expression levels of genes of interest were normalized using the primers (forward; reverse) (AGTGCCAGCCTCGTCCCGTA; TGAGCCCTTCCACAATGCCA), for expression of GAPDH (see Supplemental Data 3 for obtained values). All other primer sequences are detailed in (Liddelow *et al*., 2017; Clarke *et al*., 2018). Data were analyzed by one-way ANOVA followed by Dunnett’s multiple post hoc test.

### 2.12 Statistical analysis and cell counting

All statistical analyses were done using GraphPad Prism 8.00 software. The results are shown as mean ± SD. Statistical analysis was performed using one-way or two-way ANOVA, unless otherwise stated. Probability values of 0.05 (p<0.05) were considered to indicate statistical significance. N=biological replicates, n=technical replicates.

## 3 Results

### 3.1 OECs secrete anti-inflammatory factor(s) that reduce astrocyte reactivity

Nuclear translocation of the pro-inflammatory protein NFκB was used as a readout of astrocyte reactivity evoked by bacterial endotoxin LPS (Rothhammer *et al*., 2016), as measured by immunoblot analysis of the nuclear fraction of astrocyte lysates (Figure 1A, inset). As expected, NFκB increased in the nuclear fraction of immortalized C8D30 astrocytes treated with LPS (Figure 1A, gray bars, p≤0.05). Co-culture of astrocytes with OECs blocked nuclear translocation of NFκB in response to LPS, as previously reported (Hale *et al*., 2011; Figure 1A, purple bars). Adding CM from untreated OEC monocultures, (OEC-CM), also decreased NFκB translocation into nuclei of astrocytes exposed to LPS (Figure 1A, red bars,), indicating that anti-inflammatory factor(s) are secreted by OECs even in the absence of a stress signal.

To facilitate identification of OEC factors of interest, the anti-inflammatory capacity of six immortalized OEC lines (Calof & Guevara, 1993) were screened. Immortalized astrocytes (C8D30) were treated with CM from the different OEC lines, treated with LPS, immunostained for NFκB (Figure 1B), and the nuclear/cytoplasmic ratio of NFκB immunostaining was determined. As shown in Figure 1C, CM from two of the immortalized OEC lines, C7 and D6, significantly reduced nuclear NFκB translocation in C8D30 astrocytes compared to LPS treatment alone (Figure 1C, red bars, p<0.05), while D4 and C4 CM were similar to LPS alone. Original characterization of these immortalized OEC lines had been based on morphology and immunostainining with markers expressed by primary OECs (Calof & Guevara, 1993). Characterization of the re-grown lines was consistent with earlier reports, with C7 cells, for example, showing heterogeneous morphologies depending on culture conditions and density (Figure 1D): these included Schwann Cell spindle-like (majority; Figure 1Da), astrocyte-like type1 (Figure 1Db), and astrocyte-like type2 (Figure 1Dc) morphologies (Huang *et al*., 2008). Cell lines were re-examined by immunofluorescence for expression of OEC-specific markers, such as p75, Sox10, and brain lipid-binding protein (BLBP). Both C7 and C4 cell lines were positive for these OEC markers (Figure 1E). Since C7-CM significantly suppressed nuclear NFκB translocation in astrocytes, whereas C4-CM did not (Figure 1B, C), and both lines expressed the OEC markers tested, C7-CM was used as a positive control, and C4-CM as a negative control in further experiments.

### 3.2 OEC-secreted CryAB suppresses LPS-induced astrocyte reactivity

To identify OEC-derived molecules potentially involved in crosstalk between OECs and astrocytes, secreted proteins from primary OEC-CM, C7-CM and C4-CM were compared by mass spectrometry. Before collection of CM, LPS was added to cultures as a stress signal. Secreted proteins from LPS-treated cells were ranked based on 1) their abundance in C7-CM compared to C4-CM; 2) their abundance in C7-CM compared to primary OEC-CM; and 3) absence from C4-CM (Figure 2A). Proteins that were secreted at similar levels by C7 cells and primary OECs (Figure 2A, horizontal dashed line), but are not likely to be present in C4-CM (Figure 2A, X axis) were determined. Based on these criteria, we identified two proteins of particular interest: the heat shock protein alpha crystallin B chain (CryAB), and the cell surface glycoprotein MUC18 (MCAM). To identify OEC secreted molecules in response to an endogenous signal from astrocytes, similar experiments were performed after treatment with Anosmin1, an extracellular binding protein secreted by mature astrocytes (Gianola *et al*., 2009) and shown to act on OECs (Hu *et al*., 2019). Even though the ortholog is yet to be identified for this protein in mice, we observed a robust migration of primary mouse OECs towards recombinant Anosmin-1 (personal observation). Notably, both CryAB and MCAM were identified as major secreted proteins in this screen as well (Figure 2B). CryAB was selected for further study because of its known role as an anti-inflammatory protein involved in stress responses by CNS glia (e.g., Ousman *et al*., 2007; Kuipers *et al*, 2017), and because it was the most abundant protein fitting our criteria in screens of both LPS (Fig. 2A) and Anosmin-1 (Fig. 2B) treated samples. Recombinant CryAB protein mimicked the effect of OEC-CM or C7-CM on astrocyte reactivity, as measured by suppressed nuclear translocation of NFκB, following either LPS- or cytokine-induced inflammation (Figure 2C).

### 3.3 Exosomes secreted by OECs contain CryAB, which moderates intercellular immune response

Since it has been shown that CryAB secretion can occur via exosomes (Sreekumar *et al*., 2010; Kore *et al*., 2014; Guo *et al*., 2019), exosomes were isolated from OECs to determine whether they were positive for CryAB and whether the CryAB secreted via OEC-exosomes had the ability to attenuate astrocyte reactivity. For these experiments, exosome fractions were isolated from culture supernatants of OECs generated from both *CryAB^−/−^* mice and WT (*CryAB*^+/+^) controls. To ensure the quality of fractions used, exosomes and whole cell lysates (CL) from WT OECs were analyzed by immunoblotting for the following proteins: the structural protein, ß-actin; a mitochondrial protein, Tomm20; a nuclear protein, histone H3; and the extravesicular protein Flotilin-1. The exosome fraction was devoid of ß-actin, Tomm20 and histone H3, but was positive for Flotilin-1 (Figure 3A), consistent with published information for exosome fractions (Jeppesen *et al*., 2019). Competitive ELISA against CryAB confirmed the presence of CryAB in OEC exosomes. Exosomes derived from 1×10^6^ OECs contained 9.22 ± 0.2 ng CryAB, whereas CryAB protein was undetectable in CM from which exosomes were depleted (Supplemental Figure 1A). Next, exosomes from both genotypes were immunoblotted for CryAB and the endocytosis protein, Alix, which is concentrated in exosomes (Figure 3B; Jeppesen *et al*., 2019). CryAB was present in WT OECexo fractions but was absent in *CryAB*^−/−^OECexo fractions; while the exosome marker Alix was present in exosome fractions from OECs of both genotypes. Finally, to assay whether CryAB present in OEC exosomes could suppress astrocyte reactivity, the exosomes were added to immortalized C8D30 astrocytes for 24 hrs, and treated with LPS for the last 2 hours of this incubation. As shown in Figure 3C, quantitative immunoblotting demonstrated that: a) exosomes from WT OECs were able to suppress astrocyte reactivity, as measured by reduced nuclear translocation of NFκB; b) astrocytes treated with *CryAB*^−/−^ OECexo remained reactive; and c) the reactivity of astrocytes treated with *CryAB*^−/−^ OECexo was reduced by the presence of recombinant CryAB protein. Immunostaining OEC-astrocyte co-cultures for NFκB further showed strong nuclear NFκB immunostaining exhibited by astrocytes co-cultured with *CryAB*^−/−^ OECs (Figure 3D, lower right). whereas astrocytes cocultured with WT OECs showed little if any nuclear NFκB immunostaining (Figure 3D, lower left). Together, these results are consistent with CryAB, secreted by OECs in exosomes being an important protein for OEC-astrocyte crosstalk, and functioning as an anti-inflammatory molecule for astrocytes.

**Figure 3:**
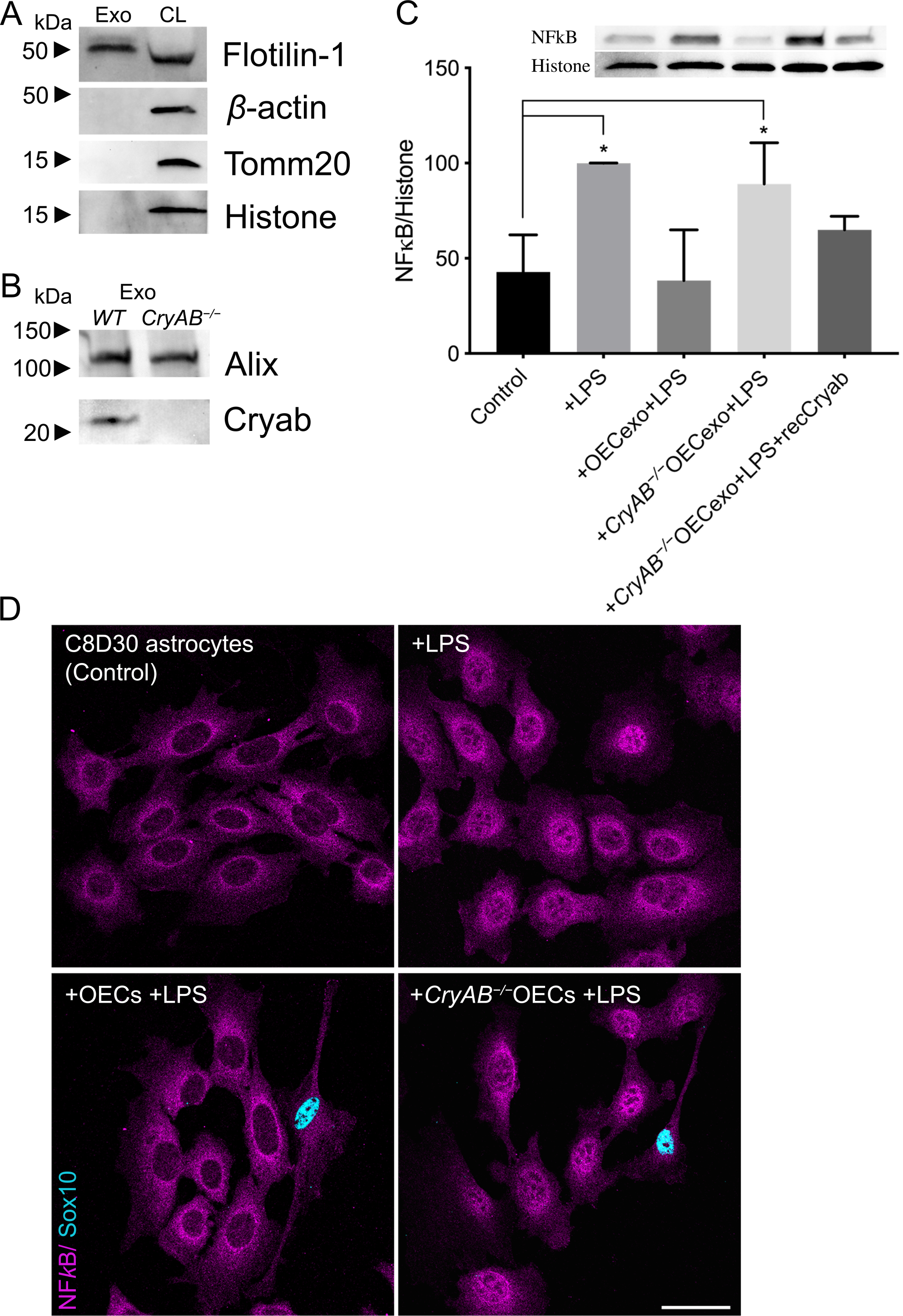
CryAB, secreted by primary OECs into exosomes, suppresses inflammatory response in an astrocyte cell line. (A) Immunoblot of exosome (Exo, left) and whole cell lysate (CL, right) fractions from WT primary OECs were screened for β-actin, Tomm20, histone H3, and Flotilin-1. The exosome fraction from WT exosomes was devoid of cellular ß-actin, Tomm20 and histone H3, but contained the extravesicular protein Flotilin-1. (B) Immunoblots for CryAB and the exosome marker Alix were performed on exosome fractions made from *CryAB*^−/−^ and WT (*CryAB*^+/+^) OEC cultures. CryAB was absent in exosome fractions from *CryAB*^−/−^ OEC culture medium, whereas the exosome marker Alix was present. (C) C8D30 astrocytes were treated for 24 hours with exosomes isolated from WT or *CryAB*^−/−^ OECs. Astrocytes were exposed to 1μg/ml LPS for the last 2 hrs of exosome treatment. Nuclear fractions of astrocytes were analyzed via quantitative immunoblotting for NFκB and Histone H3. Inset shows a representative immunoblot and graph shows mean ± SD of NFκB/histone ratio. All conditions were compared to astrocyte alone group (Control, N = 3; p ≤ 0.05; oneway ANOVA). Treatment of astrocytes with WT OEC-exosomes (exo) + LPS, blocked nuclear NFκB translocation. In contrast, (*CryAB*^−/−^ OEC-exosomes failed to suppress nuclear NFκB translocation. Recombinant CryAB (50ng/ml) added to *CryAB*^−/−^ OEC-exosomes was sufficient to attenuate NFκB translocation induced by LPS, with levels comparable to WT OEC-exo +LPS. (D) C8D30 astrocytes treated with LPS for 2 hrs (top right: “+ LPS”) showed stronger immunostaining for NFκB in the nucleus (magenta) compared to untreated controls (top left). Astrocytes co-cultured with *CryAB*^−/−^OECs (Sox 10-positive cells with blue nuclei) had increased levels of NFκB immunostaining in the nucleus (bottom right: “+*CryAB*^−/−^OECs +LPS”) compared to astrocytes co-cultured with WT OECs (bottom left: “+OECs +LPS”). Scale bar represents 40μm.

### 3.4 Astrocytes internalize CryAB-containing OEC exosomes

To determine whether astrocytes take up CryAB-containing exosomes secreted by OECs, *CryAB*^−/−^ astrocytes were cultured with exosome fractions from OEC cultures generated from WT mice. Uptake was visualized by immunostaining of GFAP-positive astrocytes (Figure 4, magenta); colabeled with antibodies to endosome/exosome marker CD63 (red) and CryAB (green). CryAB and CD63 colocalized in *CryAB*^−/−^ astrocytes treated with exosomes for 4 hrs (Figure 4, insets). Neither untreated *CryAB*^−/−^ astrocytes (Figure 4B) nor WT astrocytes (Figure 4C) showed such specific colocalization, consistent with the uptake of CryAB-containing OEC exosomes by astrocytes.

**Figure 4:**
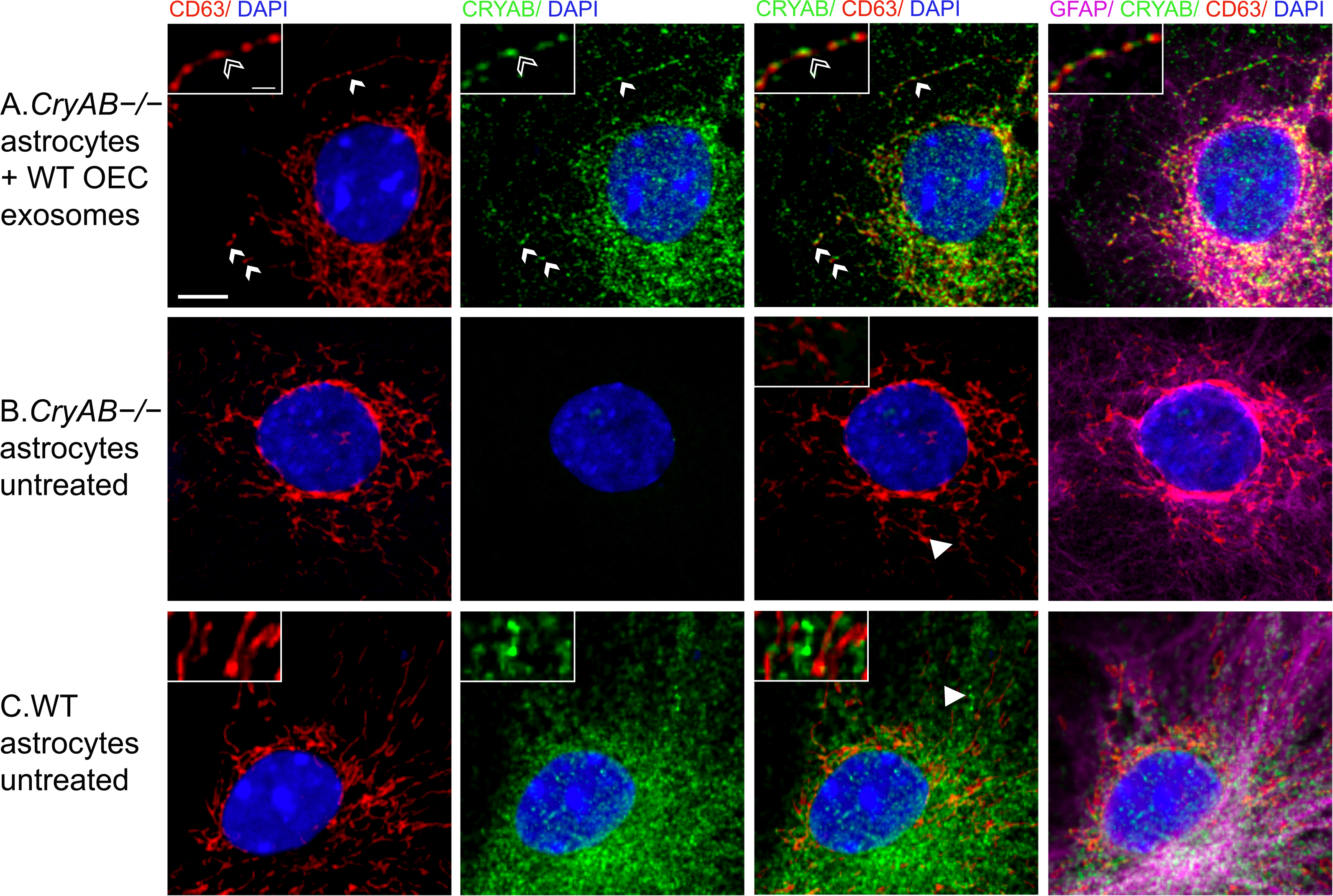
CryAB in OEC exosomes is internalized by astrocytes. (A) OEC exosomes from WT mice were co-cultured with primary astrocytes from CryAB KO mice. (B) Untreated astrocytes from CryAB KO. (C) Untreated astrocytes from WT mice. All groups were stained for CD63 (endosomes; red), CryAB (green), GFAP (magenta) and Dapi (blue). Uptake of OEC secreted CryAB (green) is detected in CryAB^−/−^ astrocytes and is often associated with endosomes (red) (A, arrows, top arrow area shown in inset, arrowhead points to CryAB positive endosome). No CryAB staining (green) is detected in untreated astrocytes from CryAB KO (B, inset). CryAB (green) is present in untreated astrocytes from WT mice but rarely associated with endosomes (C, inset, arrowhead). Scale bar represents 5μm in low mag and 1μm in insets.

### 3.5 OEC secreted factors, including CryAB, reduce astrocytes’ expression of genes associated with neurotoxic reactivity

To evaluate the effects of OEC-secreted CryAB on expression of “neurotoxic” genes, astrocytes were exposed to LPS alone; or WT OEC-CM, *CryAB*^−/−^ OEC-CM, or CryAB immunoprecipitated from isolated OEC-exosome fractions (IP-CryAB) together with LPS. mRNA from treated astrocytes was then analyzed for 12 transcripts known to be associated with neurotoxic astrocyte reactivity (Liddelow *et al*., 2017). Q-RT-PCR analysis (Figure 5) showed that all tested transcripts were reduced in expression in the presence of WT OEC-CM, and this effect was significant for 9 of the 12 (Figure 5, second row, white arrows, p<0.05 A vs B). In contrast, 4 of the transcripts showed increased expression when treated with *CryAB*^−/−^ OEC-CM (Figure 5C, black arrows). The analysis also suggests that suppression of expression of *Ggta1, Serping1, ligp1, Gbp2* and *Amigo2* was CryAB-dependent, for the following reasons: a) suppression of expression failed to occur with *CryAB*^−/−^ OEC-CM treatment, while still taking place with IP-CryAB treatment (Figure 5, C vs D); or b) expression was upregulated in the *CryAB*^−/−^ OEC-CM group (Figure 5, C vs A). Suppression of expression of 4 genes (*H2-T23, Srgn, H2D1 and C3*) appeared to be independent of CryAB, since it still occurred in astrocytes treated with *CryAB*^−/−^ OEC-CM (Figure 5, C vs A). In contrast to either OEC-CM treatments (*CryAB*^−/−^ or WT), a significant increase in *Fbln5* was detected in astrocytes treated with IP-CryAB (Figure 5, D vs A). These results are consistent with the finding that CryAB, secreted by OECs, functions as an antiinflammatory agent for astrocytes. In addition, comparison of OEC-CM treatment to IP-CryAB for transcripts *Ugt1a1, C3* and *Fbln5* suggest that there are factors in OEC-CM, in addition to CryAB, that suppress neurotoxic astrocyte reactivity.

**Figure 5:**
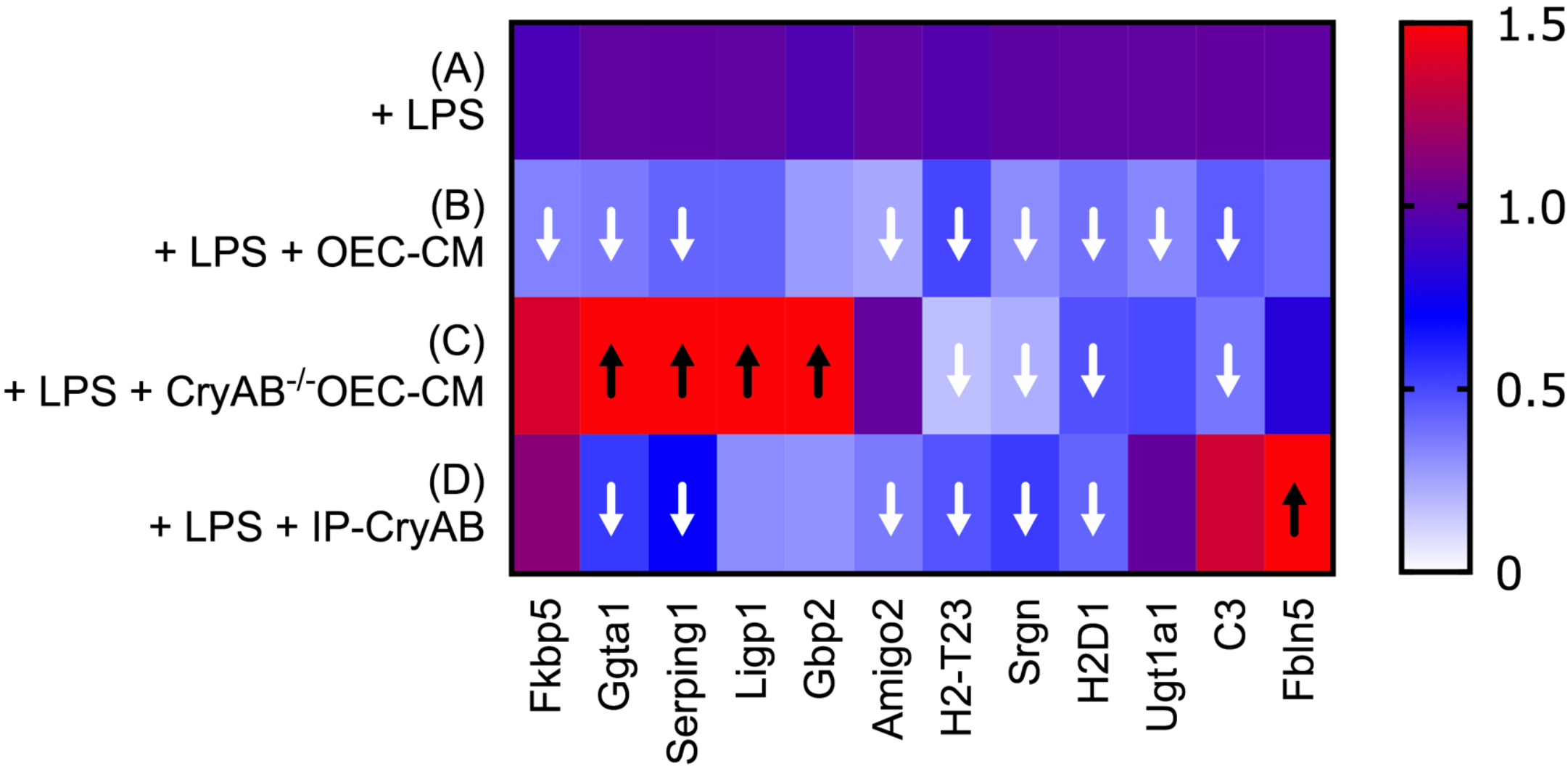
OEC-CM suppresses multiple transcripts associated with neurotoxic astrocyte reactivity in LPS-treated astrocytes. Heat map illustrating results of Q-RT-PCR (Supplemental Data 3) to detect neurotoxic astrocyte transcripts in primary astrocytes treated with LPS (A); LPS plus WT OEC-CM (B); LPS plus *CryAB*^−/−^ OEC-CM (C); or IP-CryAB for 24 hrs (D). Compared to LPS, OEC-CM (B) significantly suppressed 9 transcripts, whereas *CryAB*^−/−^ OEC-CM (C) significantly suppressed 4. (D) IP-CryAB suppressed 6 of the 9 transcripts suppressed by OEC-CM. In addition, *CryAB*^−/−^OEC-CM+LPS caused a significant increase in 4 transcripts, while IP-CryAB caused a significant increase in one. Finally, one transcript, C3, was significantly suppressed by both OEC-CM treatments (*CryAB*^−/−^ or WT) but was elevated by IP-CryAB treatment. Q-RT-PCR experiments were analyzed by one-way ANOVA followed by Dunnett’s multiple post hoc test (n= 3; arrows p ≤ 0.05).

## 4. Discussion

### 4.1 OEC-secreted factors that moderate astrocyte reactivity

Astrocyte reactivity is a pathological response that occurs in a wide range of CNS injuries, inflammation and diseases. *In vivo* studies show that some reactive astrocytes induced by ischemia can promote neural recovery and repair (reviewed in Rossi *et al*., 2007); in contrast, reactive astrocytes induced by bacterial endotoxins such as LPS are neurotoxic (Zamanian *et al*., 2012). This harmful, neurotoxic astrocyte reactivity appears to be driven by pro-inflammatory cytokines secreted by activated microglia (Liddelow *et al*., 2017). However, anti-inflammatory factors that suppress neurotoxic astrocyte reactivity are largely unknown. Olfactory system is one of the few niches in the mammalian CNS that supports neuronal regeneration (Forni *et al*., 2013). Olfactory sensory neurons are vulnerable to damage due to their exposed location in the nasal cavity and have a remarkable capacity for regeneration (Calof *et al*., 1996, Forni *et al*., 2013), suggesting the presence of robust anti-inflammatory factors in the olfactory system. OECs wrap and guide axons of the olfactory sensory neurons en route to the OB, where they establish new connections. Moreover, OECs directly interact with astrocytes at the entry point into the CNS, enabling regenerating olfactory axons to make new connections (Williams *et al*., 2004; Li *et al*., 2005; Raisman & Li, 2007). Notably, both transplanted OECs (Lakatos *et al*, 2003; reviewed in Roet & Verhaagen, 2014) and co-cultured OECs (Hale *et al*., 2011) have been shown to intermingle with astrocytes and to moderate astrocyte activation.

The studies in this report investigate anti-inflammatory factors secreted by OECs participating in OEC-astrocyte crosstalk. Mass spectrometry was used to analyze proteins in CM from primary OECs and compared to the CM of immortalized OEC lines with different anti-inflammatory capacities. Two proteins were identified as potential factors that could suppress neurotoxic astrocyte reactivity: MCAM and CryAB. MCAM (also called CD146 or MUC18) is a signaling receptor that can be cleaved from the cell membrane, generating a soluble form that is associated with increased cell migration and invasion (Seftalioglu & Karakoc, 2000), and primarily has been studied in endothelial cell angiogenesis and cancer metastasis (reviewed in Dye *et al*., 2013). Studies on Multiple Sclerosis (MS) patients showed no association between MCAM expression and disease activity (Petersen *et al*., 2019). In contrast, CryAB treatment of MS patients was associated with a therapeutic outcome, downregulation of T cell proliferation and pro-inflammatory cytokine production (Quach *et al*., 2013; van Noort *et al*., 2015). In addition, CryAB has been shown to have neuroprotective and regenerative effects in neuroinflammatory animal model systems (Arac *et al*., 2011; Ousman *et al*., 2007; van Noort *et al*., 2015). Moreover, there is a correlation between glial activation and increased CryAB levels in Alexander, Alzheimer’s and Parkinson’s diseases, as well as traumatic brain injury and stroke (reviewed in Dulle & Fort, 2016). Thus, we investigated the role of OEC-secreted CryAB in OEC-astrocyte crosstalk.

### 4.2 Exosomal release of CryAB

Our results show that OECs secrete CryAB via exosomes, and that these exosomes produce an intercellular anti-inflammatory effect following their uptake by astrocytes. Exosomes are small vesicles packaged inside multivesicular endosomes (MVE) and released into the extracellular matrix when MVE fuse with the plasma membrane (Jeppesen *et al*., 2019). Subsequent uptake by a neighboring cell initiates intercellular communication. Notably, released exosomes are functional components of the extracellular matrix that can be induced by stress signals but are associated with cell-cell communication rather than apoptosis (Jeppesen *et al*., 2019; Gupta *et al*., 2014). Consistent with our findings, recent studies suggest that CryAB can be secreted by glia in an autocrine manner (Kore *et al*., 2014; Guo *et al*., 2019) and play a protective role (Ousman *et al*., 2007). Although the downstream effects of CryAB in astrocytes that we demonstrate have yet to be fully explored, one important role of CryAB may be its interaction with transcription factors, including NFκB, to suppress inflammation by inhibition of their nuclear translocation (Shao *et al*., 2013; Zhang *et al*., 2015; Qiu *et al*., 2016). However, exosome-mediated regulation of the astrocytic immune response by OECs may also be a unique interaction, as membrane composition and protein content of exosomes is cell type (Kalra *et al*., 2012; Keerthikumar *et al*., 2015; Kim *et al*., 2015) and context specific (György *et al*., 2011; Müller *et al*., 2012). The robust anti-inflammatory response induced by OEC-secreted CryAB, shown in the present report, may also be a function of concentration, as our results indicate the concentration of CryAB in OEC exosomes to be higher than that found in astrocyte exosomes. We found that CryAB in OEC secreted exosomes was ~21% higher than that of astrocyte exosomes (Supplemental Figure 1A) and exposure to LPS increased OEC secreted CryAB concentration by ~25% (Supplemental Figure 1B), indicating OECs actively respond to stress by either secreting more exosomes or increasing CryAB concentration in exosomes, or both. In addition, exposure to Anosmin1 increased CryAB concentration by ~77% (Supplemental Figure 1C), compared to control protein recombinant myoglobin, suggesting OECs can actively respond to extracellular astrocyte signals by increasing CryAB concentrations.

### 4.3 CryAB, as well as other factors secreted by OECs, can suppress neurotoxic astrocyte reactivity

Stress causes denaturation of correctly folded proteins that can result in their aggregation and binding to CryAB (Muranova *et al*., 2018) and subsequent changes in gene expression (Singh *et al*., 2019). Our results show that OEC-secreted CryAB suppressed expression of a number of genes associated with neurotoxic astrocyte reactivity and suggest there are additional factor(s) in OEC-CM that further suppress this harmful reactivity. In fact, complement component-3 (C3) expression in IP-CryAB +LPS-treated astrocytes, although not significantly different than that observed in astrocytes treated with LPS alone, was significantly greater than expression in astrocytes treated with either OEC-CM groups (Figure 5). C3 is an important marker for neurotoxic astrocytes, evident by knockout mice showing reduced activity in microglia and astrocytes, as well as neuron loss (Shi *et al*., 2017). Moreover, C3 is found colocalized with astrocyte markers in regions of neurodegeneration in human post-mortem tissue (Liddelow *et al*., 2017). Therefore, more experiments are required to identify OEC-secreted factor(s) that can suppress C3. In this regard, other proteins identified in our mass spectrometric screen, showing smaller changes, may be worthwhile to evaluate. Certainly, crosstalk mechanisms between OECs and surrounding niche cells, including astrocytes, are highly complicated (Chuah *et al*., 2011). In addition to unidentified anti-inflammatory factors, absence or suppression of pro-inflammatory factors might also play a role in OEC-CM’s effect on astrocyte reactivity. For example, our mass spectrometry screening showed higher concentration of S100A4 and S100A6 proteins in C4-CM compared to pOEC-CM or C7-CM (Figure 2, Supplemental Data-1 & 2). S100 proteins are known to modulate neuroinflammation (Donato, 2001), and as such are also candidate molecules that may contribute to the observed difference in the inflammatory reactivity of astrocytes treated with C4-CM versus pOEC-CM or C7-CM.

Recent studies have transformed our perception of astrocytes from a passive structural support network for neurons, to active effectors in the regulation of synaptic transmission, neural excitability, plasticity and recovery. In this paper, we identify OEC secreted CryAB as an anti-inflammatory factor that can moderate astrocyte reactivity, suppressing both transcription of neurotoxic classified genes and nuclear translocation of pro-inflammatory factor NFκB. Improving our understanding of the crosstalk between astrocytes and OECs may inform strategies to identify other endogenous repair mechanisms that facilitate CNS repair, and consequently impact function, in the injured nervous system.

## Acknowledgements

We thank Dr. S. Kawauchi for help transferring OEmyc790 cell lines; Dr. Y. Li and the NINDS Protein/Peptide Sequencing Facility for mass spectrometry analysis; Dr. L. Dong and the NEI transgenic mouse core for providing cryopreserved sperm from *CryAB*^−/−^ mice; Dr. J. Pickel and the NIMH transgenic mouse core facility for rejuvenation of mice; Dr. V. Schram and the NINDS Light Imaging Facility for use of the confocal scanning and STED microscopy. We also thank the Amara Lab, Sibley Lab and Friedman Lab for the use equipment. Dr. Y. Shan, Dr. J. Chen and Dr. S. Inagaki for technical assistance. We are grateful to Drs E. Quinlan, S. Constantin, H. Cho, N. Whittington, P. Yuen, J. Street and M. Barzik for providing helpful advice.

This work was supported by the Intramural Research Program of the National Institutes of Health, National Institute of Neurological Disorders and Stroke (ZIA NS002824-28). The authors declare that there are no potential conflicts of interest.

---

**Supplementary Figure 1:**
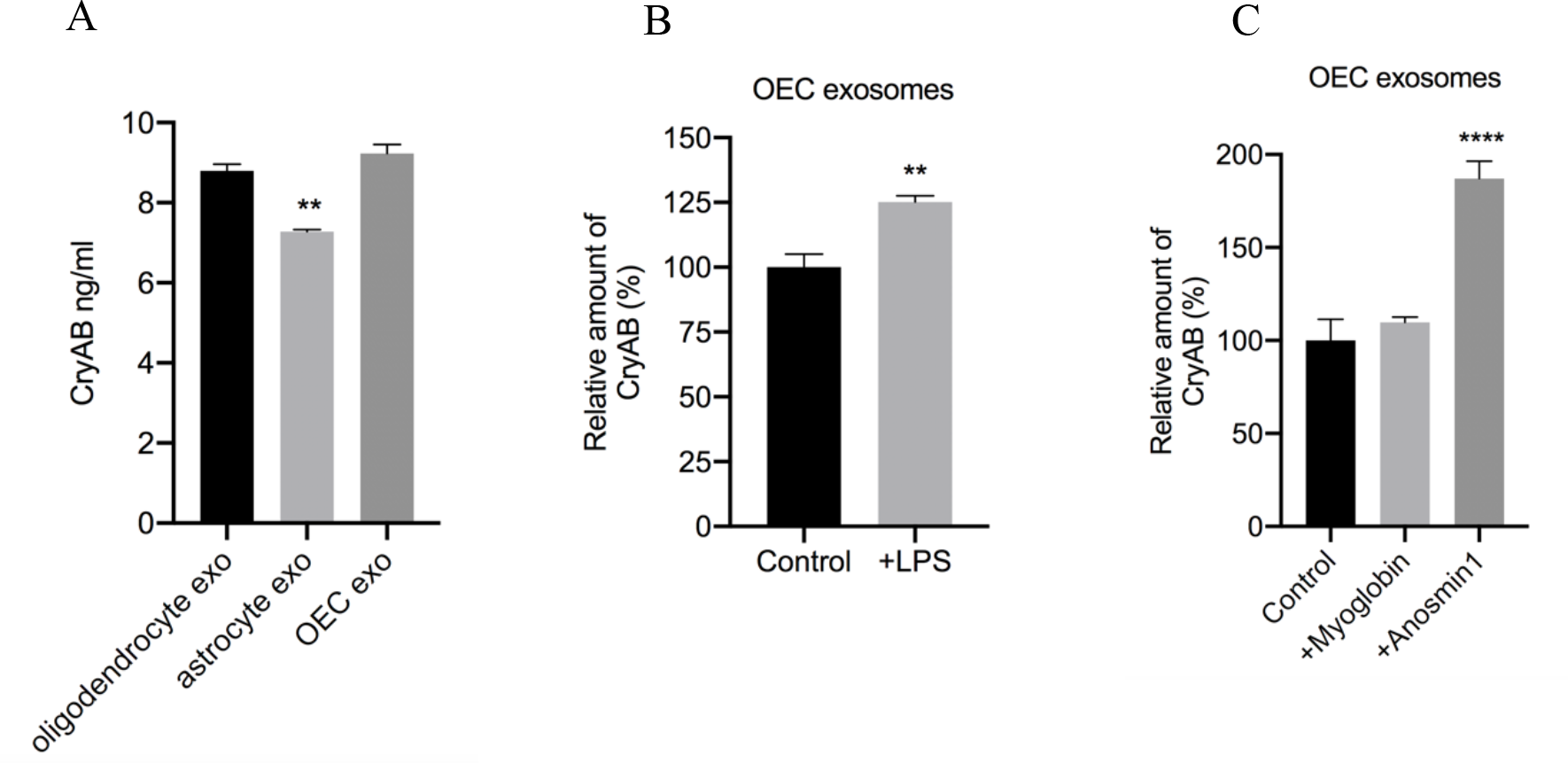
Competitive ELISA against CryAB was used to examine exosomes. (A) Exosomes (exo) derived from 1×10^6^ OECs were determined to have 9.22 ± 0.2 ng CryAB. CryAB in OEC secreted exosomes was not significantly different than the concentration in oligodendrocyte exosomes but %21.1 higher than that of astrocyte exosomes. (B) Exposure to LPS increased OEC secreted CryAB concentration by 25.17% (C). Although OECs’ exposure to LPS or astrocytic signals were not necessary for the suppressive effect of OECs on NFκB translocation in C8D30 astrocytes, whether the CryAB secretion would be facilitated by external signals was investigated by stimulating with LPS or Anosmin1. Exposure to Anosmin-1 increased CryAB concentration by 77.17%, compared to control protein recombinant myoglobin.

## SUPPLEMENTARY ONLINE MATERIAL

**SUPLLEMENTARY TABLE 1.**
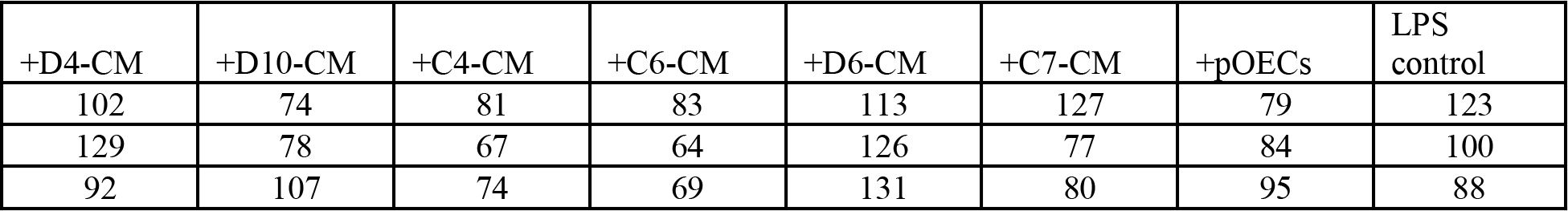

**SUPLLEMENTARY TABLE 2.**
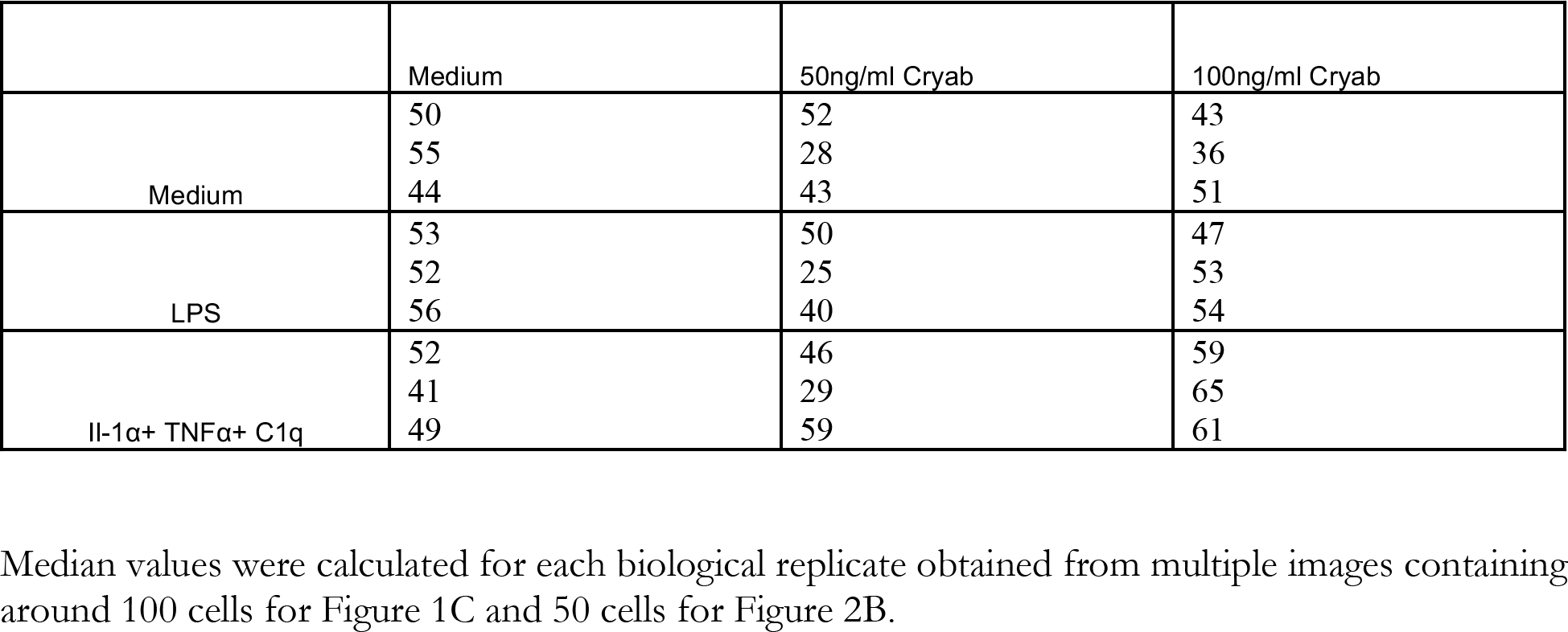

**SUPLLEMENTARY TABLE 3.**
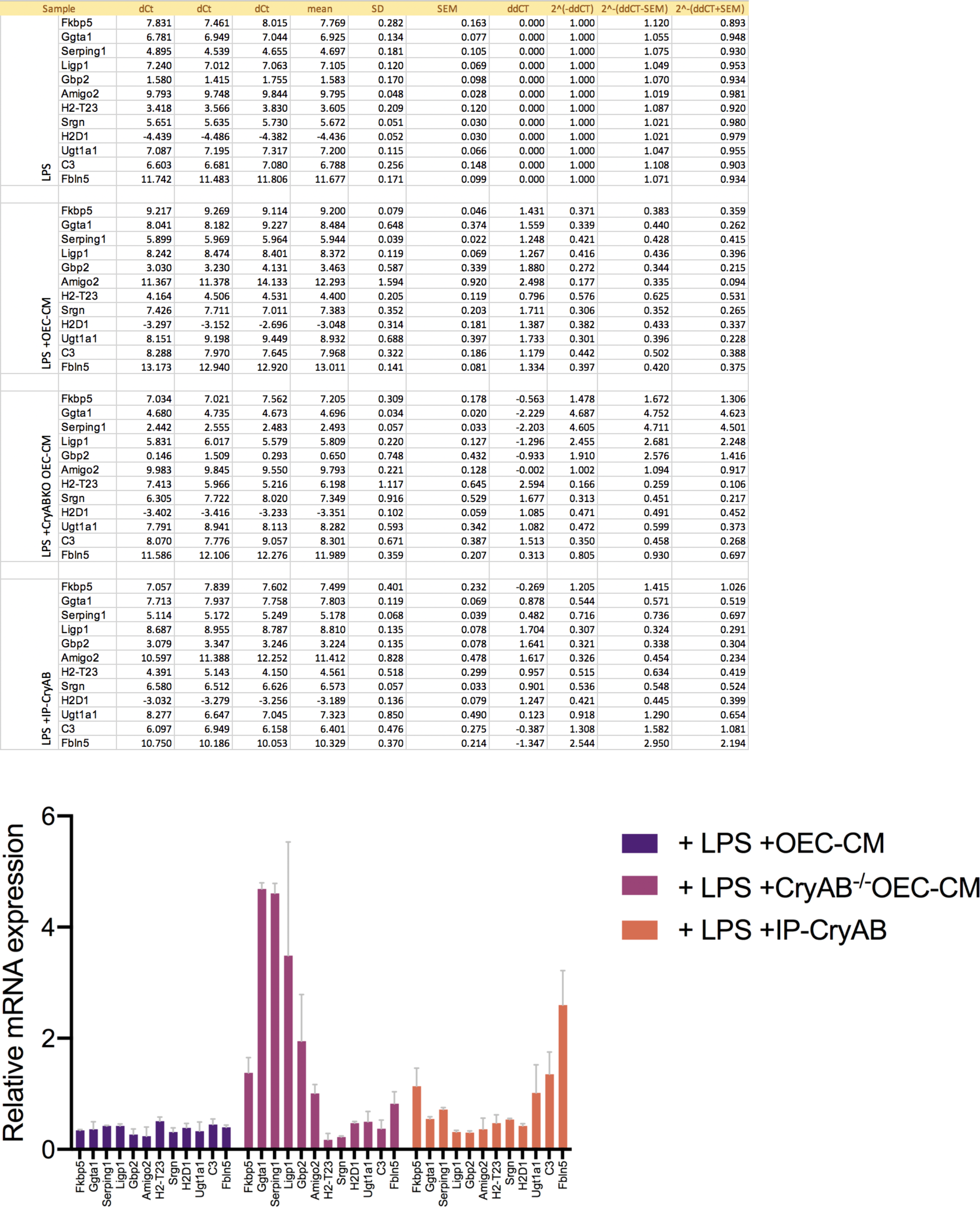
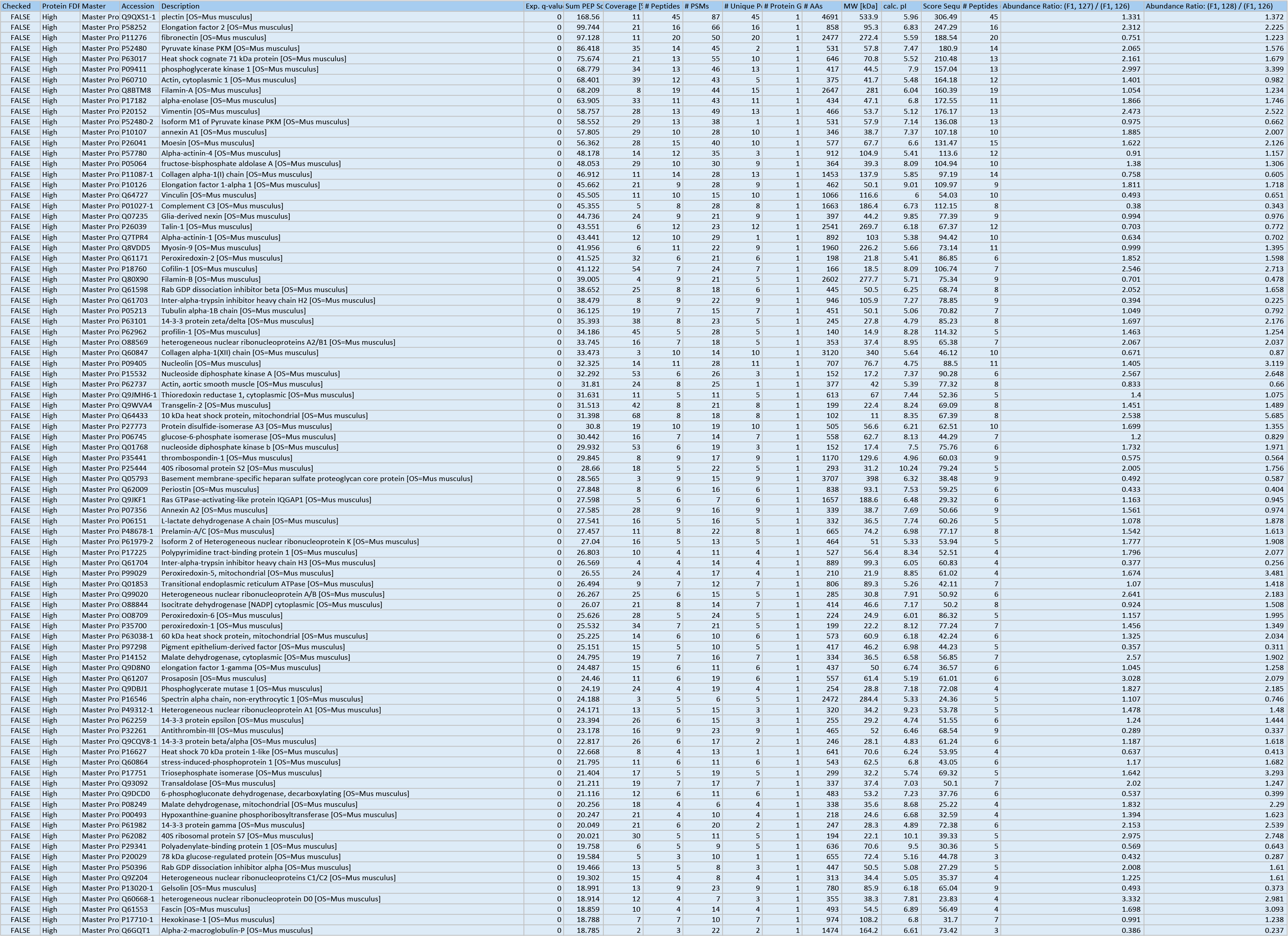
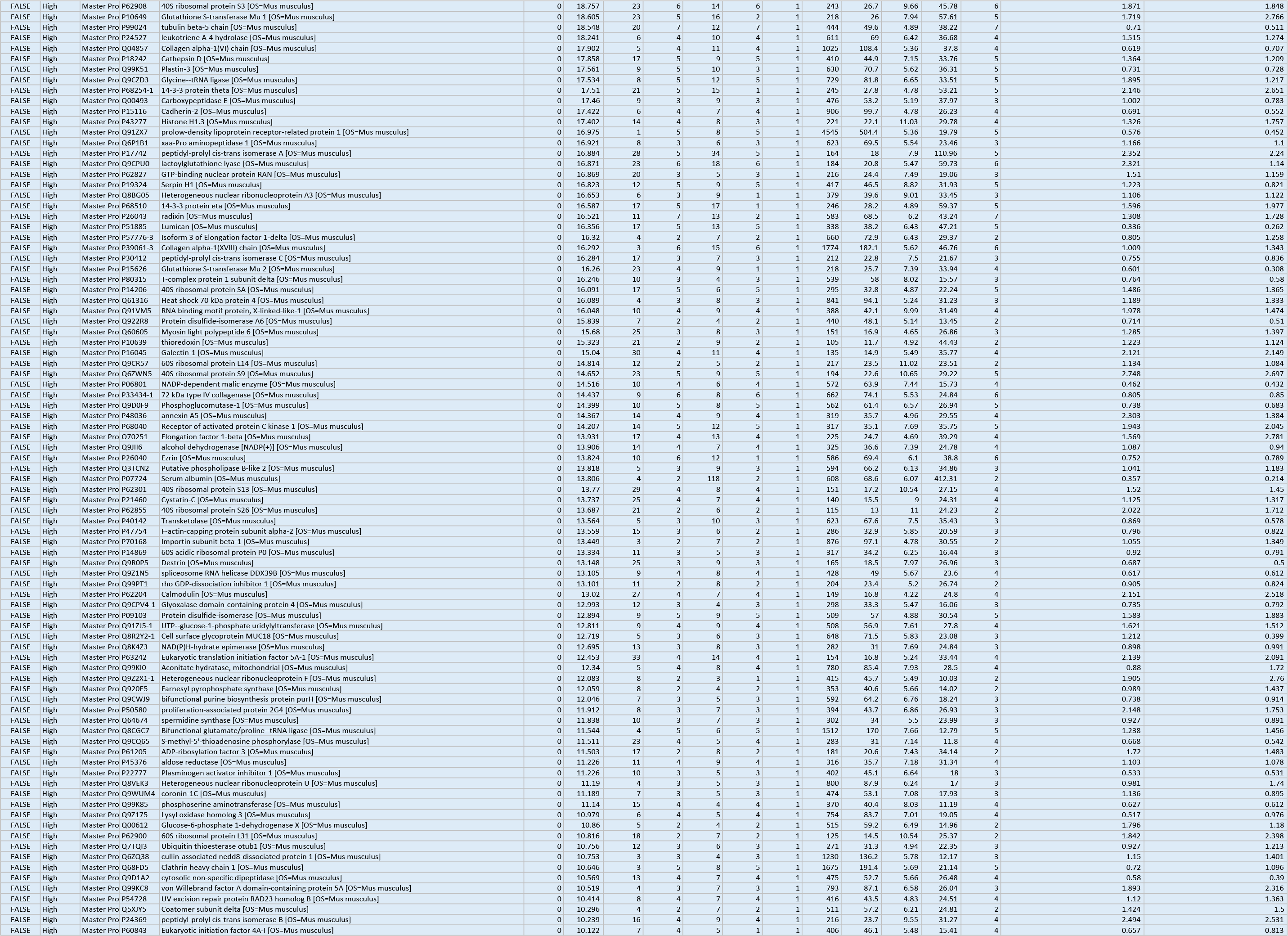
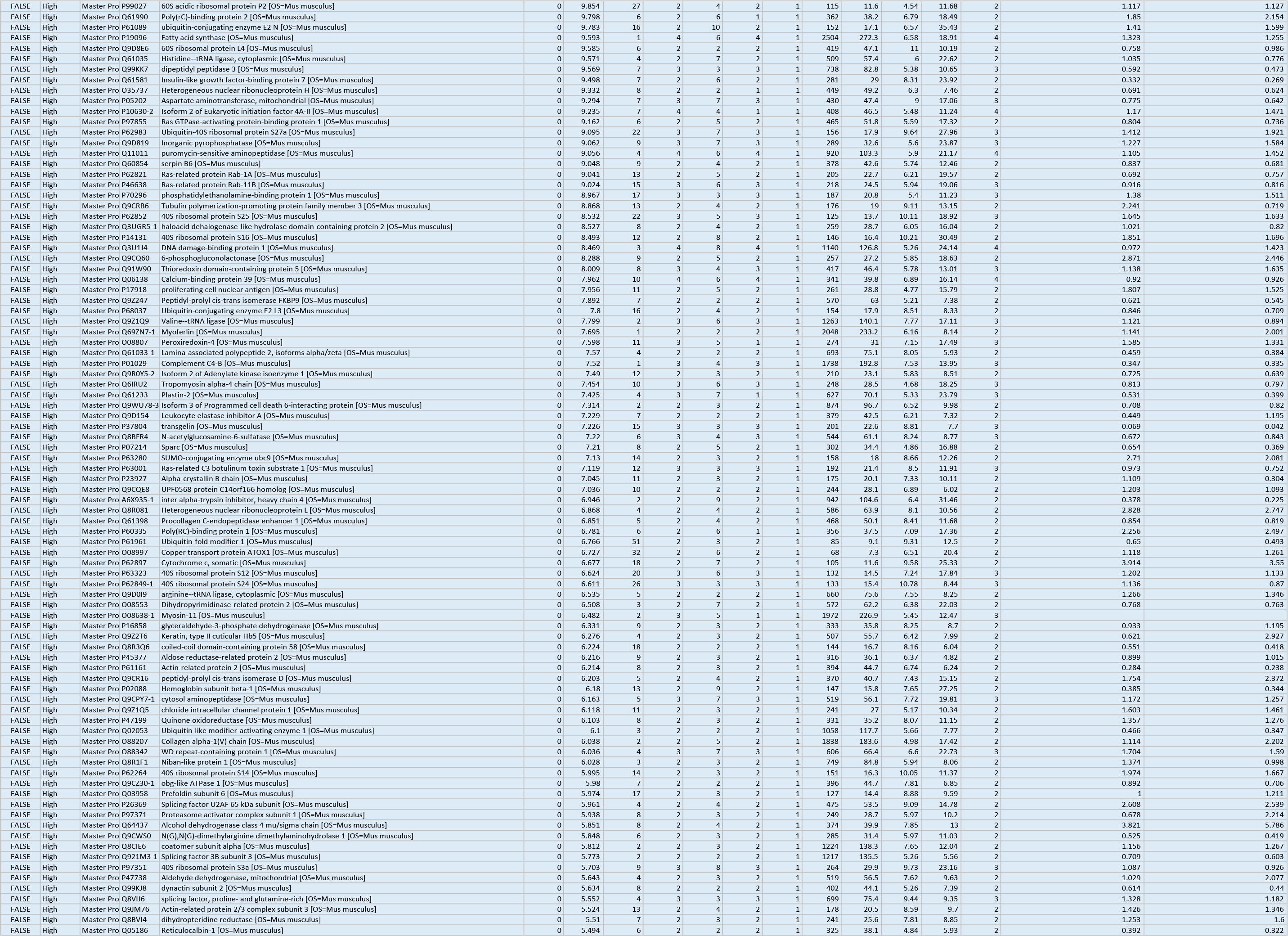
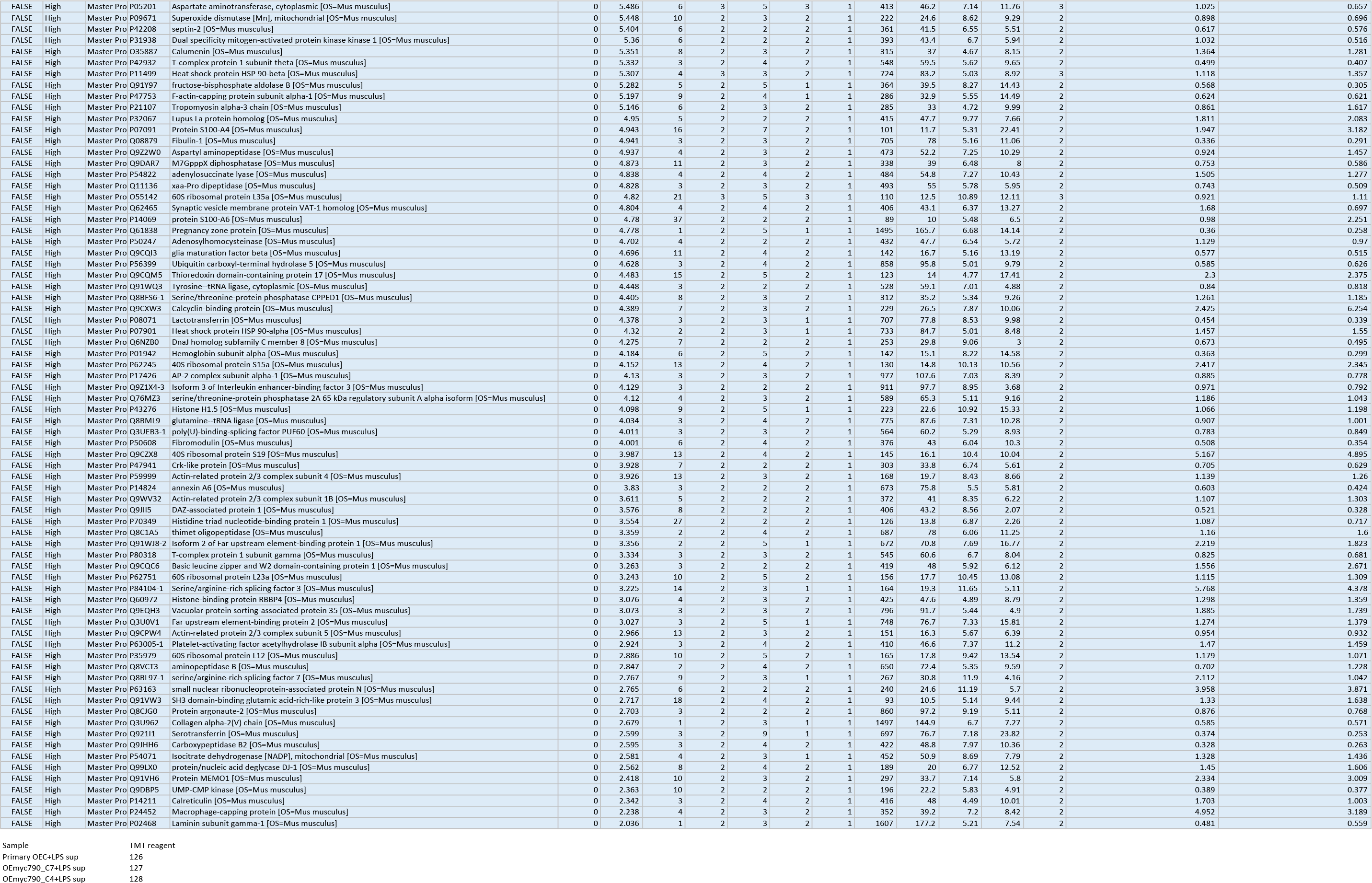
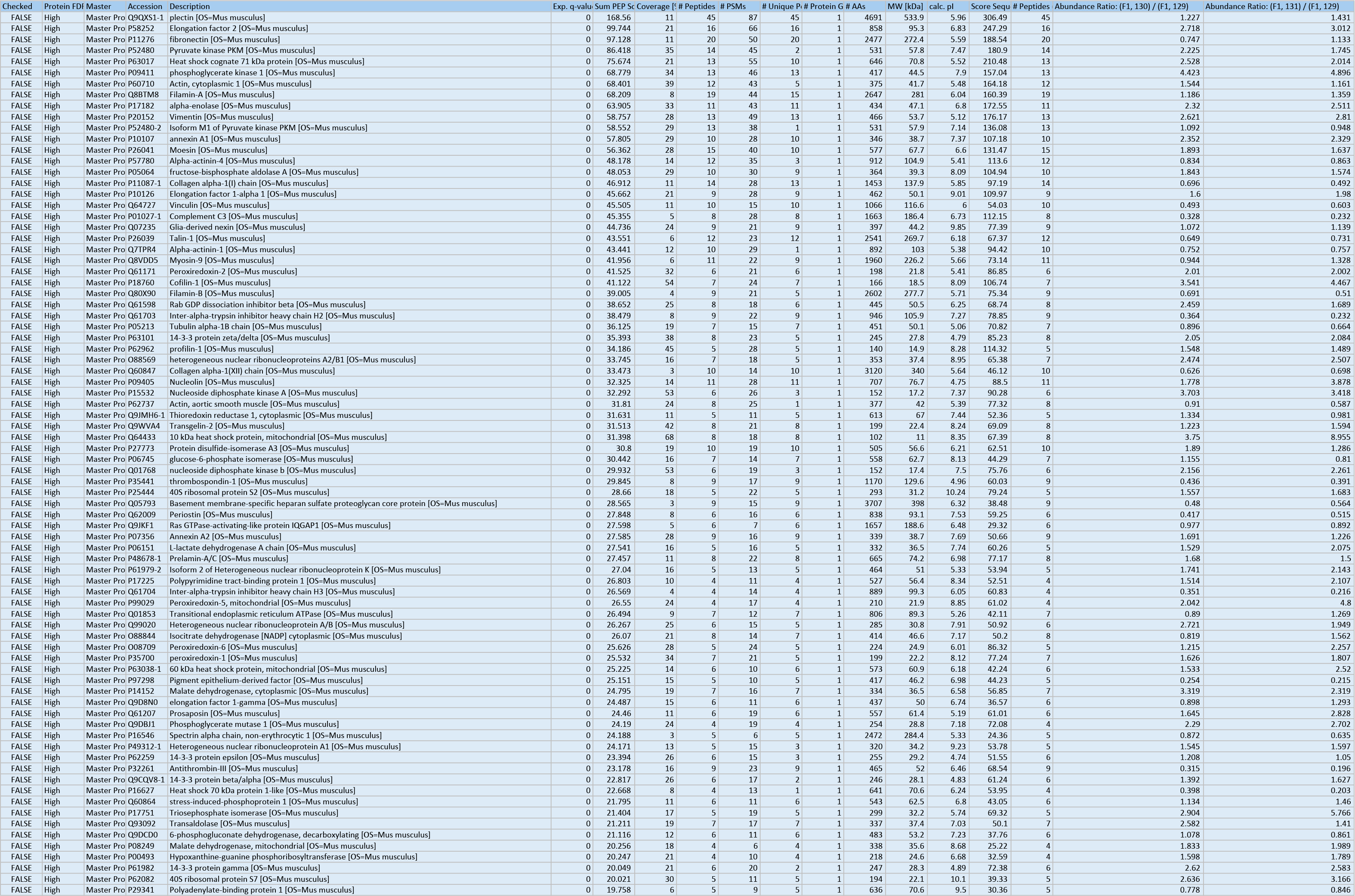
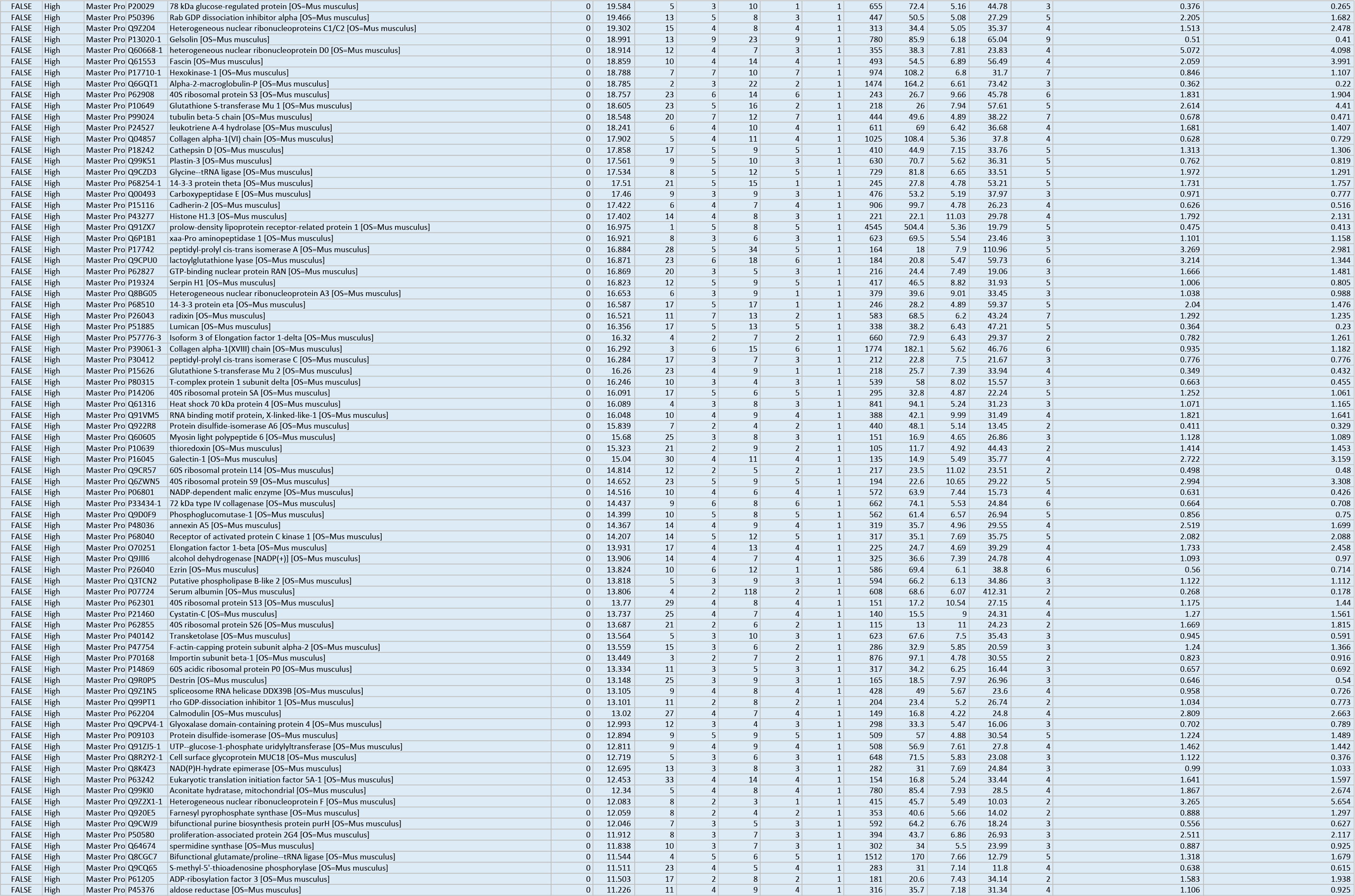
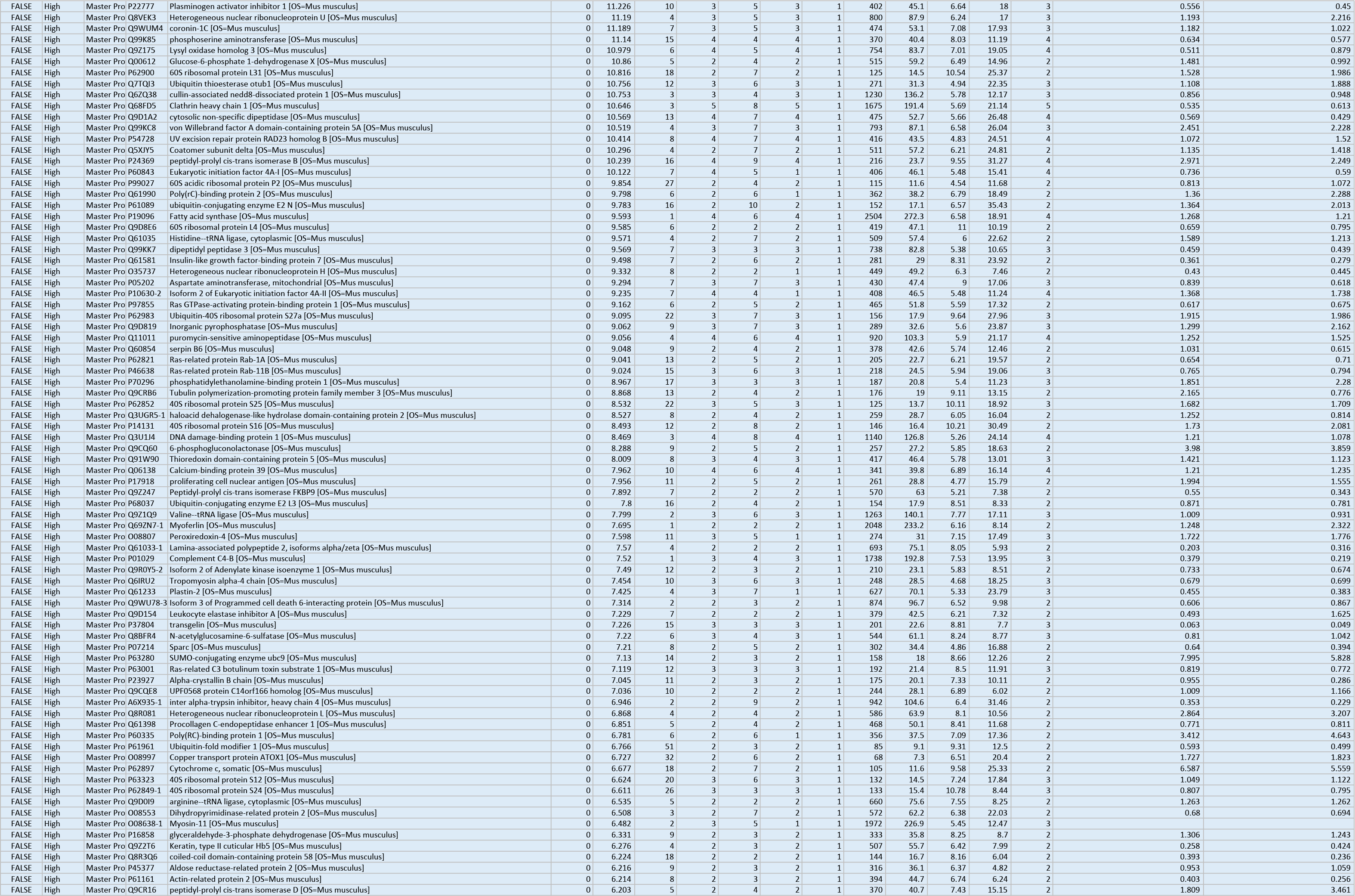
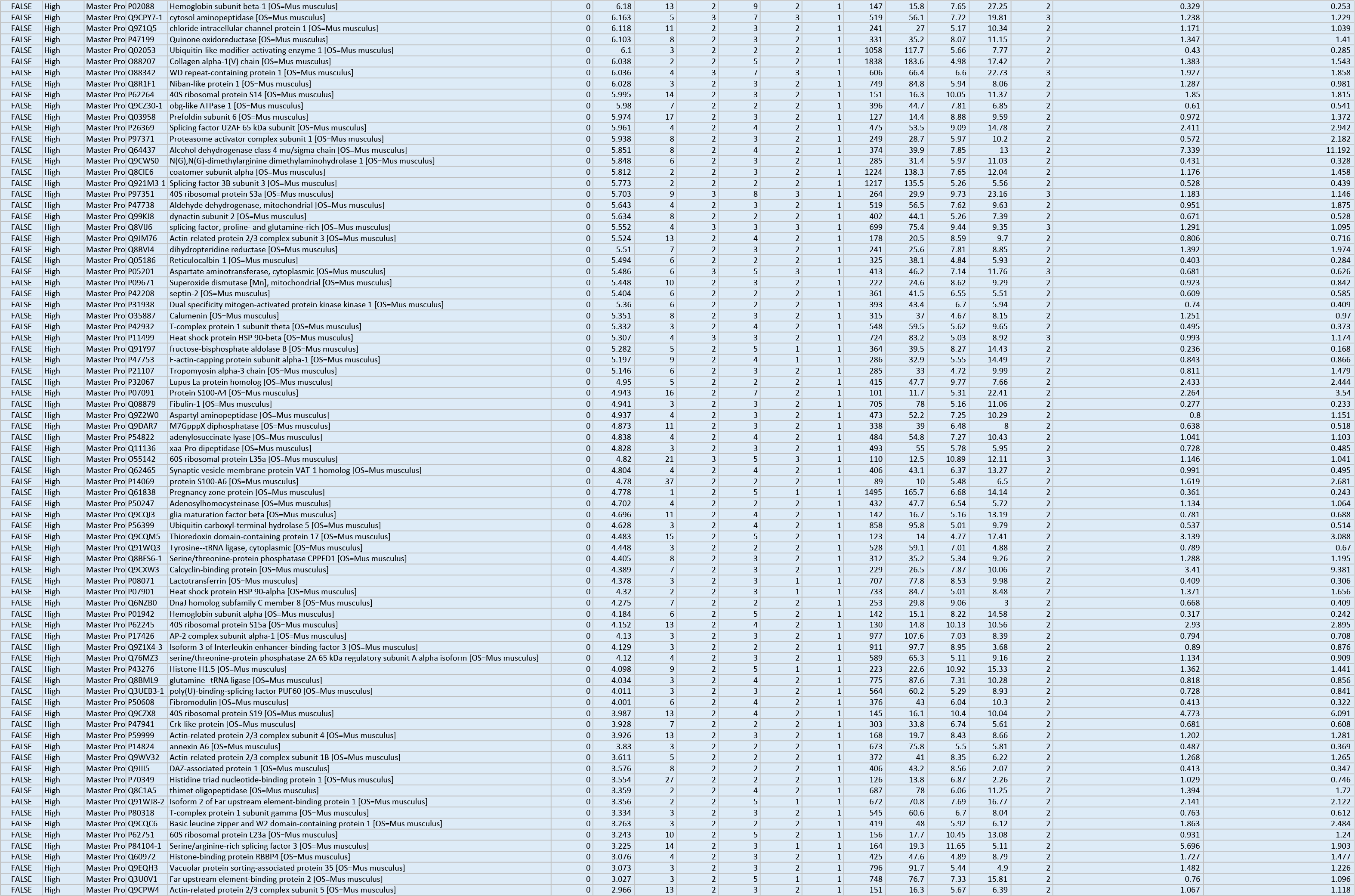
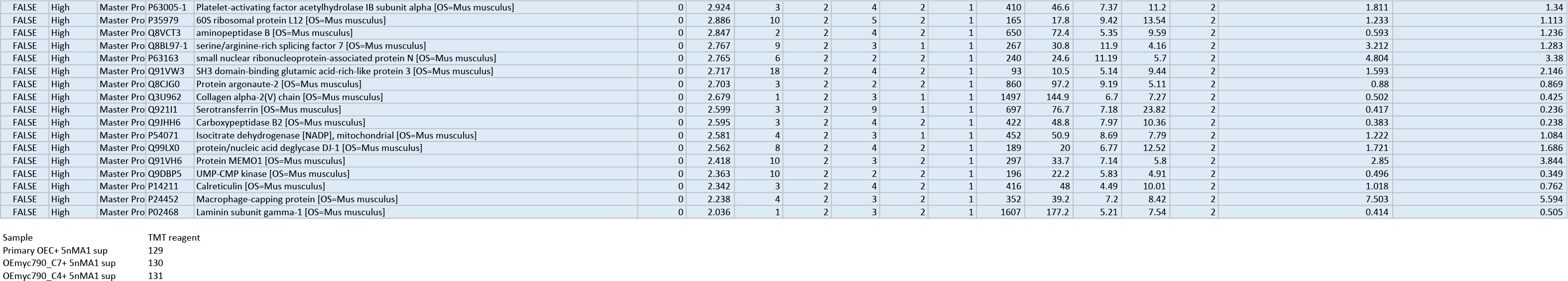

